# Uncoupling of STING Accumulation and Activation upon Chromosome-Derived Extrachromosomal Circular DNA Formation

**DOI:** 10.64898/2026.10.23.761201

**Authors:** Herve Dushimyineza, Jiawei Cai, Makoto T Hayashi

## Abstract

The innate immune system maintains cellular homeostasis by recognizing cytosolic nucleic acids. While the cGAS-STING pathway is well characterized in detecting non-self-DNA, its response to self-DNA originating from the host nucleus remains unclear. Extrachromosomal circular DNA (eccDNA), a form of self-DNA often enriched in cancer, has been implicated in immune modulation, yet its ability to activate cGAS-STING remains unresolved. Here, we developed a Cre-loxP-based eccDNA reporter system, where Cre-mediated eccDNA formation triggers expression of the fluorescent protein mCitrine, enabling time-resolved, single-cell analysis of chromosome-derived eccDNA dynamics and STING activation. Using an mRuby3-STING reporter, we found that low-dose plasmid transfection induced STING activation, as evidenced by modest STING accumulation and downstream cytokine induction. Cells carrying chromosome-derived cytosolic eccDNA also showed a modest, cGAS-dependent accumulation of STING. However, this accumulation was uncoupled from detectable STING-Ser366 phosphorylation or induction of downstream cytokines. These findings indicate that chromosome-derived eccDNA does not drive detectable STING signaling under the conditions tested, and that STING accumulation and its activation are mechanistically separable steps. This work provides a foundation for understanding the immunological roles of eccDNA in genome instability and cancer.

## Introduction

The innate immune system provides an immediate line of defense against invading pathogens and tissue damage (Medzhitov, 2007), thereby playing a central role in maintaining tissue homeostasis (Medzhitov, 2021). This system relies on pattern-recognition receptors (PRRs) that detect conserved immunogenic signals derived from either pathogens or damaged host cells (Li & Wu, 2021). These signals encompass pathogen-associated molecular patterns (PAMPs) (Medzhitov & Janeway, 2002), such as microbial nucleic acids, as well as damage-associated molecular patterns (DAMPs) (Ma et al., 2024), which originate from endogenous molecules that become mislocalized or are released upon cellular stress or cell death. Together, these sensing mechanisms coordinate antimicrobial defense by triggering inflammatory signaling cascades that culminate in the production of type I interferons (IFN-I) and pro-inflammatory cytokines (Z. Chen et al., 2025; Lin et al., 2025). In response to cytosolic double-stranded DNA (dsDNA), the cyclic GMP-AMP synthase (cGAS)-stimulator of interferon genes (STING) pathway serves as a central innate immune signaling axis (Dvorkin et al., 2024; Z. Zhang & Zhang, 2025). Upon dsDNA recognition, cGAS catalyzes the synthesis of 2ʹ3ʹ-cyclic GMP-AMP (cGAMP), which functions as a second messenger to activate STING (P. Gao et al., 2013; Sun et al., 2013; Wu et al., 2013; X. Zhang et al., 2013). Following cGAMP binding, STING undergoes sequential intracellular trafficking from the endoplasmic reticulum (ER) to the ER-Golgi intermediate compartment and subsequently to the Golgi apparatus (Hopfner & Hornung, 2020), where it recruits and activates TANK-binding kinase 1 (TBK1). TBK1 then phosphorylates STING at Ser366 (pSTING-S366), and phosphorylated STING recruits interferon regulatory factor 3 (IRF3) for phosphorylation by TBK1 (Tanaka & Chen, 2012; Liu et al., 2015; C. Zhang et al., 2019). Activated IRF3 translocates to the nucleus and drives transcription of IFN-I, while STING concomitantly activates NF-κB to induce pro-inflammatory cytokines and chemokines (Hopfner & Hornung, 2020). Secreted IFN-I then acts in an autocrine and paracrine manner to induce interferon-stimulated genes (ISGs) through JAK-STAT signaling (Schoggins, 2019). STING signaling is thus gated by trafficking rather than by cGAMP binding alone: arrival at the Golgi licenses activation through palmitoylation at Cys88/91, and blocking palmitoylation abolishes the type I interferon response despite STING translocation (Mukai et al., 2016; Jeltema et al., 2023). Whether Golgi accumulation alone is sufficient for downstream signaling, however, remains unresolved.

A critical challenge for the innate immune system is to distinguish non-self-nucleic acids from those of self-origin, thereby preventing aberrant immune activation (Bartok & Hartmann, 2020). Under homeostatic conditions, self-DNA is largely sequestered within the nucleus and mitochondria, limiting its access to cytosolic DNA sensors (Hopfner & Hornung, 2020). Although cGAS localizes within the nucleus and retains some catalytic capacity there (Gentili et al., 2019), nuclear cGAS is held in a largely inactive state by tight tethering to chromatin (Volkman et al., 2019). Structural analyses of cGAS-nucleosome complexes have revealed that contacts between cGAS and the histone H2A-H2B acidic patch occlude the DNA-binding surfaces required for activation and simultaneously block cGAS dimerization, thereby preventing assembly of the catalytically active complex on chromatinized self-DNA (Boyer et al., 2020; Cao et al., 2020; Kujirai et al., 2020; Michalski et al., 2020; Pathare et al., 2020; Zhao et al., 2020). In the cytosol, mislocalized self-DNA is additionally degraded by three-prime repair exonuclease 1 (TREX1), limiting its immunostimulatory potential (Stetson et al., 2008). Consistent with the importance of these safeguards, disruption of any of these mechanisms has been associated with chronic inflammation and autoimmune disease (Stetson et al., 2008; D. Gao et al., 2015; Dai et al., 2019). Various forms of cellular stress, including DNA damage, replication stress, and mitotic errors (Mackenzie et al., 2017), can drive the formation of micronuclei (MN), small extranuclear DNA-containing bodies, and promote the aberrant cytosolic accumulation of self-derived DNA, thereby enabling potential recognition by cytosolic DNA sensors (Mackenzie et al., 2017; M. Chen et al., 2022; Zannini et al., 2024). MN were initially proposed as activators of cGAS-STING signaling following nuclear envelope rupture and exposure of their DNA contents to the cytosol (Bartsch et al., 2017; Harding et al., 2017; Mackenzie et al., 2017). However, a growing body of single-cell-level evidence indicates that MN are not potent activators of the cGAS-STING pathway. Using a sister chromatid fusion reporter combined with a STING reporter, we showed that MN do not induce STING activation or downstream interferon responses (Sato & Hayashi, 2024). Consistently, cGAS localization to MN is uncoupled from downstream cGAS-STING activation across multiple genotoxic stress conditions (Takaki et al., 2024). Further corroborating these findings, both an intracellular cGAMP reporter showed that genotoxic-stress-induced cGAS activation is rare and is not driven by enrichment at micronuclei (Lebrec et al., 2026) and a STING-IRF3 interaction biosensor independently confirmed that ruptured MN do not activate STING signaling (Smarduch et al., 2025). Collectively, these convergent lines of evidence suggest that MN-associated chromatin DNA is largely immunologically inert, raising the broader question of whether chromosome-derived self-DNA can intrinsically activate the cGAS-STING pathway in living cells and, if not, at which step signaling is restricted.

One candidate source of immunostimulatory DNA is extrachromosomal circular DNA (eccDNA), circular dsDNA molecules that exist independently of chromosomes (Turner et al., 2017; Yang et al., 2022). eccDNA is frequently detected in tumor cells (Verhaak et al., 2019) and is thought to arise from genomic instability events, including breakage-fusion-bridge (BFB) cycles and chromothripsis, in which catastrophic chromosomal rearrangements generate DNA fragments that undergo aberrant repair and randomly circularize (Shoshani et al., 2021). Beyond its established roles in oncogene amplification, intratumoral heterogeneity, and therapy resistance (Shimizu, 2021; Zheng et al., 2025), eccDNA has recently attracted attention as a potential innate immune activator: purified eccDNA acts as a potent innate immunostimulant in a manner dependent on its circularity and on STING, independently of sequence (Wang et al., 2021). Nevertheless, it remains largely unresolved whether nuclear chromosome-derived eccDNA, which may be associated with inhibitory chromatin as a form of self-DNA, can intrinsically activate the cGAS-STING pathway in living cells.

Addressing this question is complicated by the limitations of existing eccDNA induction strategies. Most experimental approaches have relied on inducing DNA damage or cellular stress to promote eccDNA formation. Early studies linked eccDNA formation to DNA-repair-associated processes: eccDNA generation was linked to transcriptional activity and the mismatch repair pathway, possibly arising from replication slippage followed by mismatch repair (Dillon et al., 2015), and chromosome-derived eccDNA was found to be common in healthy human somatic tissue, implicating endogenous mutational processes as a source (Møller et al., 2018). More direct experimental induction of eccDNA has been achieved through genotoxic agents such as etoposide and UV irradiation, which trigger apoptosis and the associated formation of circular DNA molecules through apoptotic DNA fragmentation and DNA ligase 3-dependent circularization (Wang et al., 2021). Fragmentation of cytoplasm-exposed chromatin in ruptured micronuclei by the endonuclease N4BP2 has similarly been shown to initiate chromothripsis and ecDNA formation in cancer models (Krupina et al., 2025). However, these approaches are coupled to mitochondrial perturbation through multiple mechanisms: genotoxic insults trigger voltage-dependent anion channel (VDAC) oligomer-dependent release of mitochondrial DNA (mtDNA) into the cytosol (Kim et al., 2019; Guan et al., 2023; Kim et al., 2023; Sato & Hayashi, 2024), and apoptosis induction activates BAK/BAX-mediated mitochondrial outer membrane permeabilization, similarly liberating mitochondrial nucleic acids into the cytosol (McArthur et al., 2018; Tigano et al., 2021). Unlike nuclear DNA, mtDNA lacks histone protection and is therefore not subject to chromatin-mediated cGAS suppression, rendering it a potent cGAS activator (West et al., 2015; Kim et al., 2023; Sato & Hayashi, 2024). Therefore, any cGAS-STING response observed in stress-based systems cannot be unambiguously attributed to eccDNA itself, underscoring the need for a controlled eccDNA induction system that does not perturb mitochondrial integrity.

To address this limitation, we adapted the Cre-loxP recombination system, widely employed for conditional genomic excision (Sauer & Henderson, 1989), to develop an inducible eccDNA reporter system. A Cre-inducible strategy has recently been used to engineer large oncogene-containing ecDNAs and to examine their contribution to tumorigenesis in cells and in mice (Pradella et al., 2025). Our system is designed for a different purpose: it generates circular DNA from a genome-integrated, chromatinized reporter sequence, while simultaneously allowing single-cell-level, time-resolved tracking of eccDNA formation through a fluorescent reporter. By separating eccDNA production from mitochondrial confounders, this approach provides a well-controlled platform for interrogating the intrinsic capacity of chromosome-derived eccDNA to activate the cGAS-STING pathway in individual living cells. Combined with an mRuby3-STING reporter that monitors STING accumulation at the Golgi (Sato & Hayashi, 2024) and with readouts of STING activation, defined here by Ser366 phosphorylation and downstream gene induction, this system allows each step of the pathway to be assessed separately.

## Results

### Development of a Cre-loxP-dependent eccDNA reporter system

Our Cre-loxP eccDNA reporter is based on a DNA cassette in which two exons encoding the N- and C-terminal fragments of the mCitrine protein are arranged such that full-length mCitrine is expressed only upon Cre-dependent circularization of the loxP-flanked sequence (Fig. 1A). A neomycin resistance gene (neoR), included downstream of the loxP site, enables selection of cells carrying the integrated reporter, but is not included in the circularized product, which allows the eccDNA to be distinguished from the genome-integrated reporter (Fig. 1A). The entire cassette was integrated into the host genome via the Sleeping Beauty (SB) transposon system (Ivics et al., 1997; Turchiano et al., 2014).

**Figure 1.**
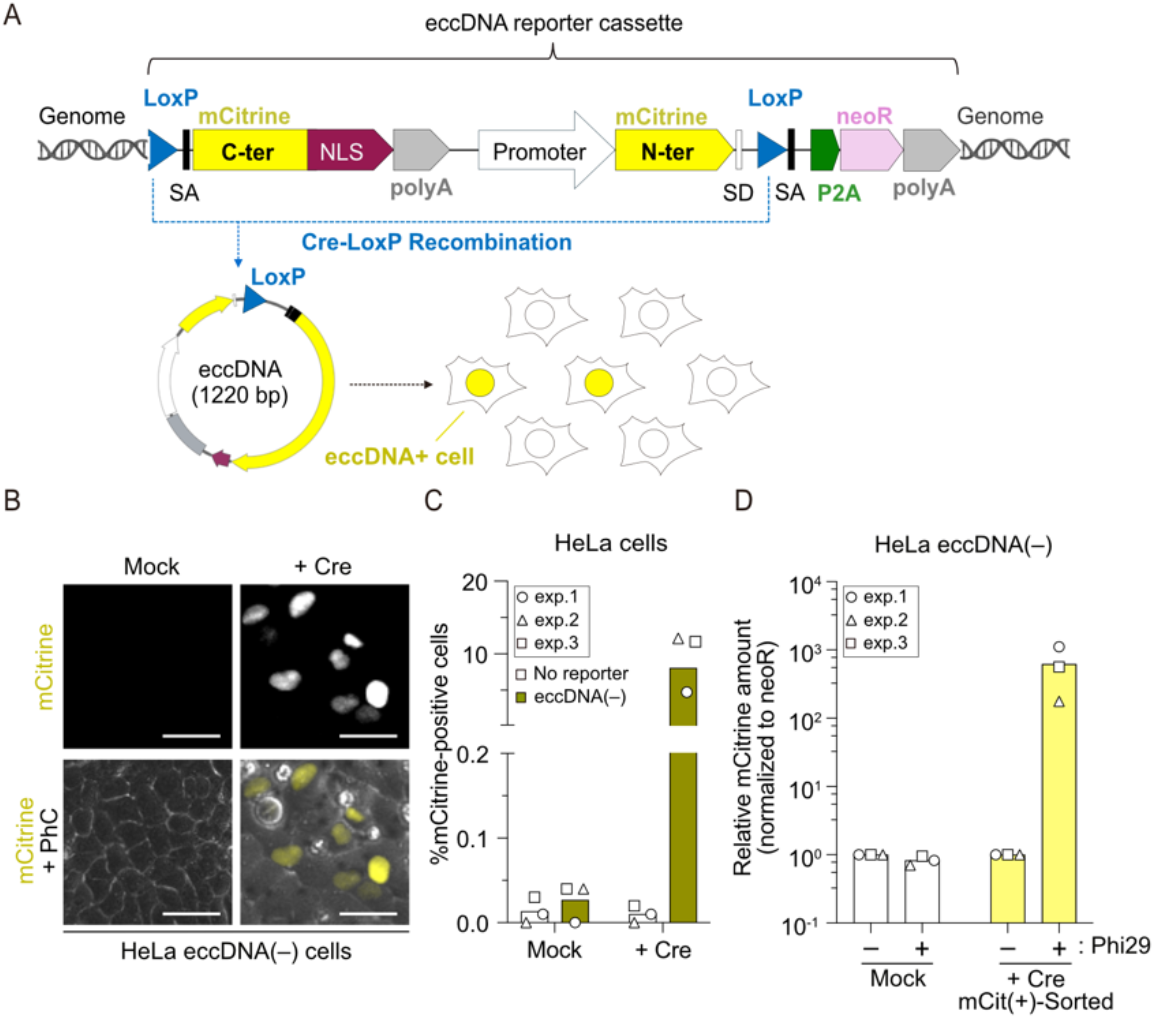
Cre-mediated cassette excision provides a defined eccDNA reporter system. **(A)** Schematic of the Cre-loxP-dependent eccDNA reporter system. Cre-mediated excision of the loxP-flanked cassette generates eccDNA and induces mCitrine expression. SA, splice acceptor; SD, splice donor; P2A, porcine teschovirus-1 2A self-cleaving peptide; NLS, nuclear localization signal. **(B)** Representative live-cell images of mCitrine in HeLa eccDNA(-) reporter cells 2 days after Cre transduction. Mock-treated cells served as controls. PhC, phase contrast. Scale bars: 25 μm. **(C)** Percentage of mCitrine-positive HeLa no reporter and eccDNA(-) reporter cells 2 days after Cre transduction. **(D)** Relative amount of mCitrine sequence following rolling circle amplification (RCA) with or without Phi29 DNA polymerase. qPCR signals were normalized to neoR and the corresponding Phi29-negative control. **Data information: (C, D)** Bars represent the mean of three independent biological replicates.

HeLa1.2.11 cells were co-transfected with two plasmids encoding the SB transposase and the eccDNA reporter donor cassette and selected with G418. Stable integrants were obtained without detectable mCitrine expression. The resulting cells, hereafter referred to as HeLa eccDNA(-) cells, were subsequently transduced with a lentivirus encoding Cre recombinase. mCitrine fluorescence was observed only after Cre transduction, indicating Cre-dependent recombination of the reporter (Fig. 1B, C). To validate eccDNA formation, we employed rolling circle amplification (RCA) using Phi29 DNA polymerase, which preferentially amplifies circular DNA templates under isothermal conditions (Dean et al., 2001; Diaz-Lara et al., 2016).

Following Cre transduction, mCitrine-positive HeLa eccDNA(-) cells were isolated by fluorescence-activated cell sorting (FACS), with mCitrine-negative cells sorted from mock-treated cultures serving as a negative control (Fig. S1A, B). Genomic DNA extracted from these sorted populations was subjected to RCA, and the products were analyzed by quantitative PCR (qPCR) to quantify mCitrine and neoR sequences (Fig. S1A). Consistent with the design prediction (Fig. 1A), the mCitrine sequence was strongly enriched relative to neoR only in the RCA products from mCitrine-positive cells (Fig. 1D), indicating that mCitrine expression reflects the presence of circular DNA molecules. The same reporter cassette also drove mCitrine expression comparable to that in HeLa eccDNA(-) cells upon Cre transduction in two additional cell lines, human fibrosarcoma HT1080 and human colorectal carcinoma HCT116 (Fig. S1C, D), indicating that the reporter system functions across diverse cellular contexts. Taken together, these results establish a Cre-loxP-dependent eccDNA reporter system in which mCitrine expression depends on Cre-mediated DNA circularization, providing a platform for investigating chromosome-derived eccDNA in living cells.

### eccDNA reporter sequence and copy number influence mCitrine expression

Having validated the reporter, we next introduced additional sequences into the cassette to test whether sequence composition influences the cGAS-dependent innate immune response. Because telomeric regions have been implicated as hotspots for innate immune activation (Y.-A. Chen et al., 2017; Nassour et al., 2024), we inserted 114 TTAGGG repeats interrupted by 13-bp non-telomeric spacers into the cassette (Fig. S2A). As a non-telomeric control, we inserted a 691-bp length-matched subtelomeric sequence from chromosome Xp (Fig. S2A). Both sequences were initially inserted between the polyA and the EF-1α promoter (Fig. S2A), and the resulting reporters were integrated into HeLa1.2.11 cells, yielding HeLa eccDNA(XpST) and HeLa eccDNA(TEL) cells. Genomic integration of each reporter was confirmed by genomic PCR (Fig. S2B).

Flow cytometry revealed that HeLa eccDNA(XpST) displayed a higher percentage of mCitrine-positive cells than HeLa eccDNA(-) following Cre transduction, and eccDNA formation in this line was confirmed by RCA-qPCR (Fig. 2A, B). In contrast, Cre transduction induced only a very small population of mCitrine-positive cells (<1%) in HeLa eccDNA(TEL) (Fig. 2A). RCA-qPCR analysis of this line revealed that the rare mCitrine-positive cells contained detectable circular DNA (Fig. 2C), whereas Cre-transduced mCitrine-negative cells did not (Fig. 2D), suggesting that telomeric repeat insertion upstream of the EF-1α promoter impairs Cre-loxP-mediated eccDNA formation through an as yet uncharacterized mechanism.

**Figure 2.**
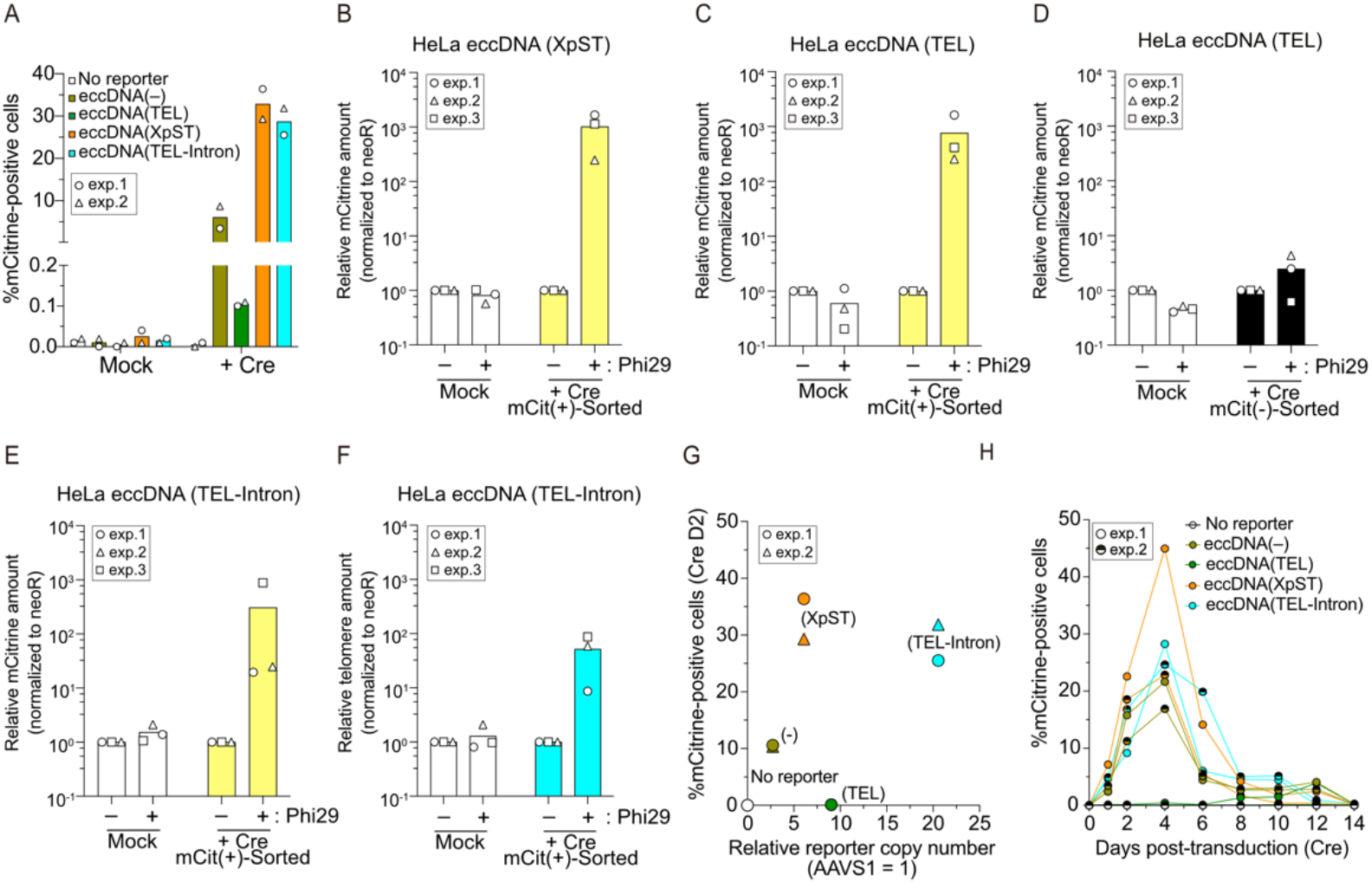
eccDNA reporter sequence and copy number influence mCitrine expression. **(A)** Percentage of mCitrine-positive HeLa no reporter, eccDNA(-), eccDNA(XpST), eccDNA(TEL), and eccDNA(TEL-Intron) reporter cells 2 days after Cre transduction. Mock-treated cells served as controls. **(B-F)** Relative sequence amount following RCA with or without Phi29 DNA polymerase. qPCR signals were normalized to neoR and the corresponding Phi29-negative control. HeLa eccDNA(XpST) cells (B), mCitrine-positive (C) and mCitrine-negative (D) HeLa eccDNA(TEL) cells, and HeLa eccDNA(TEL-Intron) cells (E, F). mCitrine and telomere sequence amounts are shown in (B-E) and (F), respectively. **(G)** Relationship between relative reporter copy number (AAVS1 = 1) and the percentage of mCitrine-positive cells at 2 days after Cre transduction in the indicated reporter cell lines. **(H)** Time-course analysis of mCitrine-positive cells in the indicated HeLa eccDNA reporter cell lines from day 1 to day 14 after Cre transduction. **Data information: (A-F)** Bars represent the mean of two independent (A) and three independent (B-F) biological replicates. **(G, H)** Data were obtained from two independent biological replicates.

To circumvent this limitation, we moved the telomere sequence into the intron downstream of mCitrine exon 1 (TEL-Intron configuration) (Fig. S2A). HeLa eccDNA(TEL-Intron) cells were established by SB-mediated integration (Fig. S2B) and expressed mCitrine upon Cre transduction (Fig. 2A). RCA-qPCR further confirmed eccDNA formation in HeLa eccDNA(TEL-Intron) cells following Cre transduction (Fig. 2E). Importantly, qPCR for telomeric repeats in the RCA products confirmed that the eccDNA derived from HeLa eccDNA(TEL-Intron) cells contains the inserted telomeric repeats (Fig. 2F).

The variability in mCitrine expression levels across reporter variants led us to examine whether the genomic copy number of the integrated cassette underlies these differences. Genomic DNA from each variant was analyzed by qPCR to determine the copy number of the mCitrine gene relative to the endogenous AAVS1 locus, using a reference plasmid (pMTH864) containing single copies of both sequences. Plotting mean copy number against the percentage of mCitrine-positive cells at day 2 post Cre-transduction revealed that mCitrine expression increased with copy number in the eccDNA(-), eccDNA(XpST), and eccDNA(TEL-Intron) reporters, whereas the eccDNA(TEL) reporter was a clear outlier (Fig. 2G and S2C). Time-course analysis further showed that HeLa eccDNA(TEL) cells maintained only a minimal fraction of mCitrine-positive cells throughout a two-week observation period, indicating sustained suppression relative to the other reporter variants (Fig. 2H). By contrast, in all other reporter variants, mCitrine expression appeared within the first few days after Cre transduction, peaked at approximately day 4, and declined from day 6 onward (Fig. 2H). This transient expression pattern is consistent with the known properties of eccDNA: owing to the absence of centromeres, eccDNA undergoes unequal partitioning during mitosis and is progressively diluted across cell divisions in the absence of selection for its retention (Shimizu et al., 1996; Turner et al., 2017; Verhaak et al., 2019).

To further dissect insert-specific behavior under defined copy-number conditions, we isolated single-cell-derived subclones from each reporter line. In subclones carrying permissive configurations, different subclones possessed variable copy numbers and mCitrine expression tended to increase with copy number, reinforcing the conclusion that eccDNA output is primarily governed by genomic dosage in these contexts (Fig. S2D-F). In contrast, eccDNA(TEL) subclones exhibited highly homogeneous copy numbers, suggesting they are derived from a few parental clones (Fig. S2G). Comparison of mCitrine expression across different reporter subclones with a relative copy number of about 5 revealed attenuated mCitrine expression in all sequence-inserted reporters relative to eccDNA(-) (Fig. S2H). Among these, eccDNA(TEL) showed the strongest attenuation, implying that the repressive effect of telomeric repeats operates independently of cassette dosage. Collectively, these findings suggest that sequence insertion into the eccDNA(-) cassette can reduce reporter output. They also indicate that copy number is the primary determinant of eccDNA reporter output in permissive configurations, whereas the position of the insert can override dosage effects. On the basis of these results, the eccDNA(XpST) and eccDNA(TEL-Intron) reporters, which support efficient eccDNA formation, were used in the subsequent experiments.

### Development of eccDNA reporter cells equipped with a STING reporter

We previously defined the STING accumulation index (St-AI) as the ratio of the maximum to the mean mRuby3-STING fluorescence intensity within individual cells, and showed in HCT116 cells that the increase in St-AI upon Compound 3 treatment reflects STING accumulation at the Golgi apparatus (Sato & Hayashi, 2024). To test whether St-AI can be used in HeLa1.2.11 cells, we transduced this line with a lentivirus encoding mRuby3-STING and isolated mRuby3-expressing cells by FACS (Fig. S3A). HeLa1.2.11 mRuby3-STING cells were then transfected with titrated doses of a GFP-encoding plasmid as a source of naked cytosolic dsDNA, and St-AI was quantified in living cells at 48 hours post-transfection. Both GFP-positive and GFP-negative cells exhibited elevated St-AI (Fig. 3A, B, and S3B-E), and the minimum plasmid dose sufficient to induce a statistically significant increase in St-AI was determined (Fig. 3B, and S3B-E). Importantly, expression of the STING-downstream chemokine *CCL5* was markedly upregulated at this same threshold dose (Fig. 3C and S3F), indicating that, in this setting, an increase in St-AI accompanies downstream gene induction by cytosolic DNA. Based on these results, we defined St-AI > 1.9 as indicating STING accumulation, a threshold chosen to maximize the difference between mock-treated and plasmid-transfected cells (Fig. 3D).Similarly, treatment with the STING agonist Compound 3 increased both the percentage of St-AI > 1.9 and *CCL5* expression at the same threshold concentration (Fig. 3E-H, and S3G, H).

**Figure 3.**
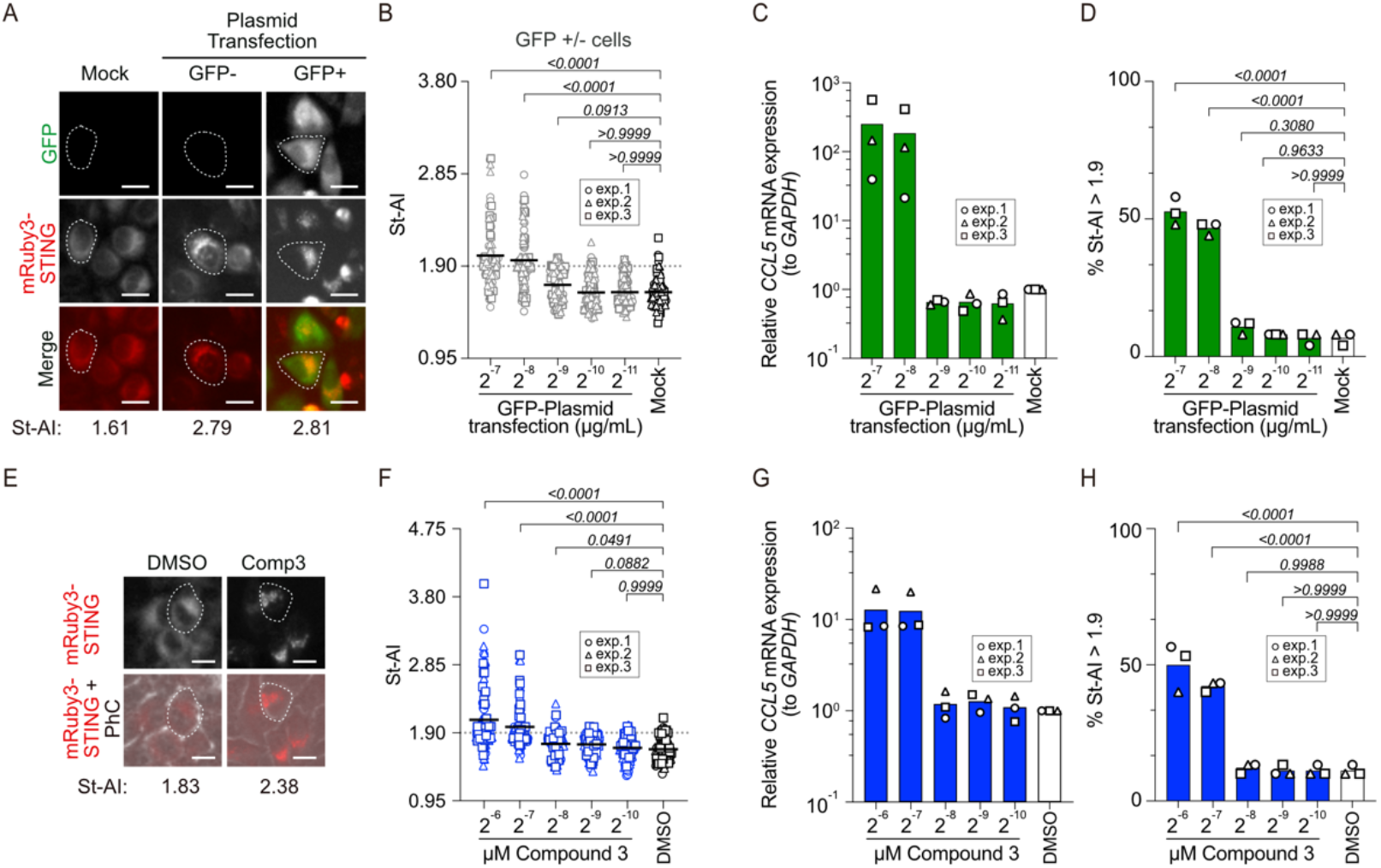
STING accumulation index reflects STING activation in response to plasmid DNA. **(A)** Representative images of GFP and mRuby3-STING in HeLa mRuby3-STING cells 48 h post-transfection with a GFP-expressing plasmid. White dotted lines indicate cell boundaries, and representative St-AI values are shown. Scale bar: 25 μm. **(B, C)** St-AI analysis (B) and RT-qPCR analysis of *CCL5* mRNA normalized to *GAPDH* (C) in HeLa mRuby3-STING cells transfected with the indicated amounts of GFP-expressing plasmid. **(D)** Percentages of cells with St-AI > 1.9 calculated from the analyses shown in (B). **(E)** Representative images of mRuby3-STING in HeLa mRuby3-STING cells treated with DMSO or Compound 3 (Comp 3, 5 h treatment followed by 19 h recovery). White dotted lines indicate cell boundaries, and representative St-AI values are shown. Scale bar: 25 μm. **(F, G)** St-AI analysis (F) and RT-qPCR analysis of *CCL5* mRNA normalized to *GAPDH* (G) in HeLa mRuby3-STING cells treated with the indicated dose of Compound 3. **(H)** Percentages of cells with St-AI > 1.9 calculated from the analyses shown in (F). **Data information: (B, F)** Bold lines represent the mean (B, n = 75 cells in total, 25 cells/biological replicate; E, n = 90 cells in total, 30 cells/biological replicate). **(C, D, G, H)** Bars represent the mean of three independent biological replicates. Statistical significance was determined by the Kruskal-Wallis test followed by Dunn’s multiple-comparison test for (B, F) and by ordinary one-way ANOVA followed by Dunnett’s multiple-comparison test for (D, H). P values are indicated for each comparison.

### A dox-inducible Cre system generates chromosome-derived eccDNA that leaks into the cytosol without detectable mtDNA release

To examine STING responses to eccDNA generation, we selected low-copy (LC, relative copy number ∼10) and high-copy (HC, relative copy number ∼100) subclones from the eccDNA(XpST) and eccDNA(TEL-Intron) reporter lines (Fig. S2E and F) and transduced these with mRuby3-STING. Because constitutive Cre expression activates STING signaling (Fig. S4A) (Pépin et al., 2016), we developed a doxycycline (dox)-inducible Cre (iCre) system to avoid this confounding effect. Time-course and dose-response analyses showed that 1 µg/mL dox treatment for 24 hours did not induce *CCL5* expression in HeLa no reporter iCre cells (Fig. S4B). Importantly, dox induced mCitrine expression in HeLa eccDNA(-) iCre cells in a dose-dependent manner (Fig. S4C), and mCitrine expression was also induced in HeLa eccDNA(XpST) and eccDNA(TEL-Intron) iCre lines (Fig. S4D). RT-qPCR analysis demonstrated that dox-induced iCre expression alone did not induce cGAS-STING downstream cytokine and chemokine genes, whereas Compound 3 treatment and plasmid transfection robustly activated these genes in HeLa no reporter iCre cells (Fig. S4E). These results indicate that the iCre system allows us to examine cGAS-STING signaling in response to eccDNA formation without confounding activation by Cre itself.

Time-course flow cytometry in eccDNA(XpST) LC/HC and eccDNA(TEL-Intron) LC/HC mRuby3-STING iCre cells revealed that mCitrine expression was first detected at 6 hours post-dox treatment and increased until 36 hours and declined by 48 hours in all subclones (Fig. 4A and B). HC subclones consistently exhibited higher mCitrine expression than their LC counterparts, as expected from copy-number-dependent reporter output. We first asked whether eccDNA accumulates in the cytosol, since nuclear-to-cytoplasmic leakage is expected to be required for activation of cytosolic DNA sensors. Cytosolic fractionation followed by qPCR revealed that the abundance of the mCitrine or telomere sequence in cytosolic fractions was markedly increased after 36 hours, but not 12 hours, of dox treatment in all reporter iCre subclones (Fig. 4C-F and S5A-C), indicating that eccDNA enters the cytosol between 12 and 36 hours of dox treatment. Transfection of each reporter subclone with the corresponding eccDNA donor cassette plasmid served as a positive control and, as expected, increased the cytosolic levels of mCitrine or telomere sequence. By contrast, the genomic AAVS1 locus, used as a control, did not accumulate in the cytosol under any condition tested (Fig. 4G, H, and S5D, E). Moreover, qPCR for the mitochondrial gene NADH dehydrogenase 1 (*MT-ND1*) detected no increase in cytosolic mitochondrial DNA under any condition tested (Fig. 4I, J, and S5F, G). Together, these results indicate that chromosome-derived eccDNA accumulates in the cytosol within 36 hours of dox treatment without detectable release of mitochondrial DNA.

**Figure 4.**
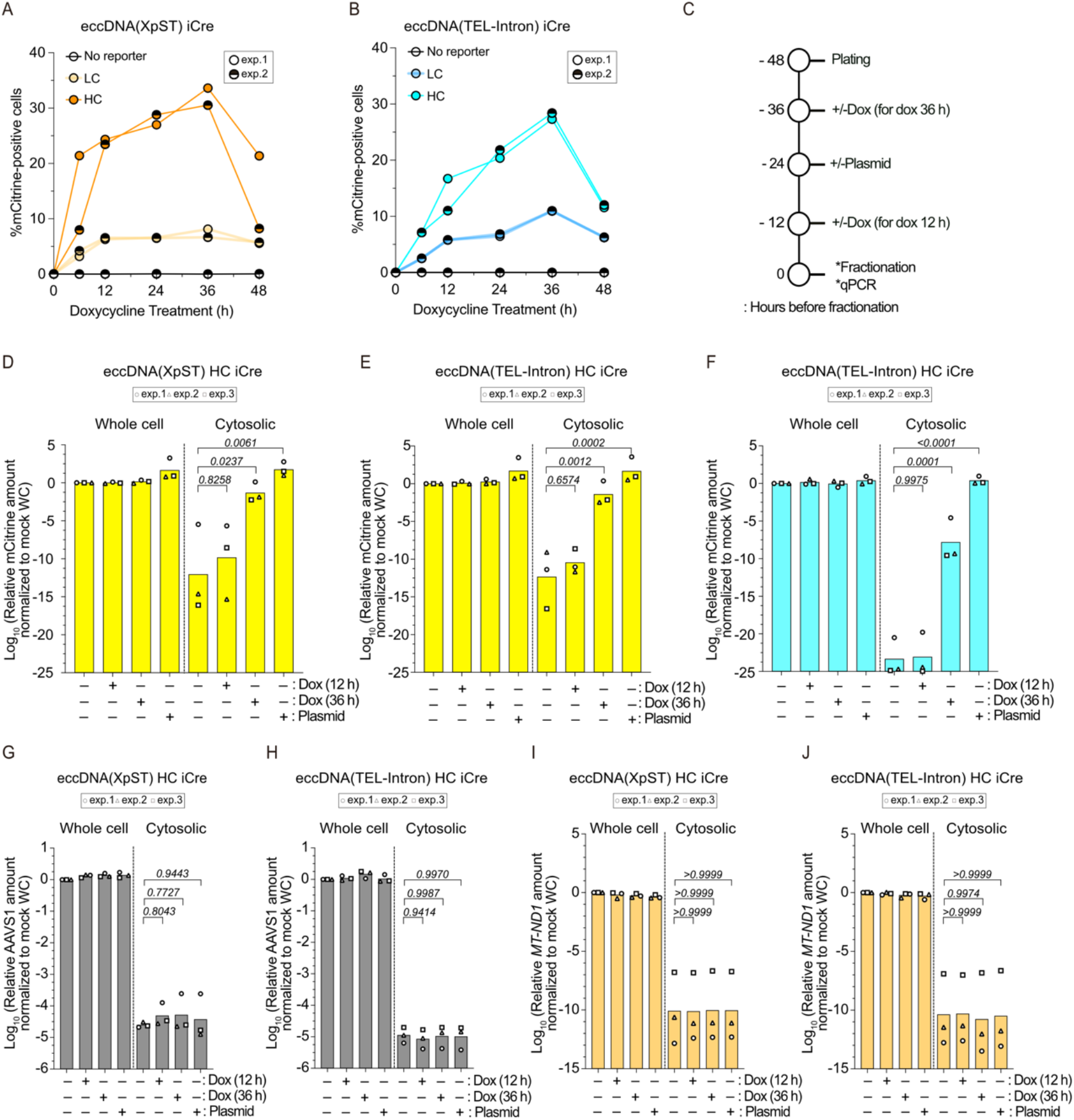
Chromosome-derived eccDNA leaks into the cytosol. **(A, B)** Time-course analysis of mCitrine-positive cells in HeLa eccDNA(XpST) mRuby3-STING iCre (A) and HeLa eccDNA(TEL-Intron) mRuby3-STING iCre (B) reporter subclones following dox treatment. LC, low copy; HC, high copy. **(C)** Experimental design for cytosolic fractionation and qPCR analysis. Cells were treated with dox for 12 or 36 h, or transfected with the eccDNA donor plasmid for 24 h, before fractionation. **(D-J)** Relative mCitrine (D, E), telomere (F), AAVS1 (G, H), and *MT-ND1* (I, J) sequence amount in whole-cell and cytosolic fractions following dox treatment or plasmid transfection in HeLa eccDNA(XpST) HC iCre cells (D, G, I) and HeLa eccDNA(TEL-Intron) HC iCre cells (E, F, H, J). qPCR signals for each target were normalized to the corresponding mock whole-cell sample and shown as log10 fold change. **Data information: (A, B)** Data were obtained from two independent biological replicates. **(D-J)** Bars represent the mean of three independent biological replicates. Statistical significance was determined by ordinary one-way ANOVA followed by Dunnett’s multiple-comparison test. P values are indicated for each comparison.

### Chromosome-derived eccDNA induces modest, cGAS-dependent STING accumulation

Based on these findings, we isolated mCitrine-positive cells from eccDNA(XpST) LC/HC and eccDNA(TEL-Intron) LC/HC mRuby3-STING iCre cultures by FACS at 12 hours post-dox treatment, before cytosolic eccDNA was detectable, and monitored St-AI daily for 3 days after sorting (Fig. 5A); days 1, 2, and 3 post-sorting correspond to 36, 60, and 84 hours post-dox, respectively. Plasmid-transfected cells showed a robust increase in St-AI on the day of sorting in all cell lines, which gradually declined thereafter, possibly reflecting progressive attenuation of STING signaling (Fig. 5B-E and S6A, B). Dox-treated HeLa no reporter iCre cells showed no change in St-AI throughout the experimental timeline, indicating that Cre expression *per se* does not activate the cGAS-STING pathway under these conditions (Fig. 5C). Similarly, no increase in St-AI was observed on the day of sorting in any of the eccDNA reporter subclones (Fig. 5D, E, and S6A, B), consistent with the absence of cytosolic eccDNA at this time point. However, St-AI showed a modest but significant increase on days 1, 2, and 3 post-sorting in all subclones, irrespective of insert sequence composition or copy number (Fig. 5D, E, and S6A, B), suggesting that cytosolic eccDNA can engage cGAS. To test this, we transduced the reporter subclones with a lentivirus encoding shcGAS-1 4 days before dox treatment or plasmid transfection (Fig. 5F and S6C). The St-AI increase induced by plasmid transfection was abolished by shcGAS-1 (Fig. 5G, H), indicating that STING accumulation induced by cytosolic DNA depends on cGAS. Similarly, the modest St-AI elevation following 36 hours of dox treatment was abrogated in shcGAS-1-expressing cells (Fig. 5G, H), indicating that eccDNA-induced STING accumulation also depends on cGAS.

**Figure 5.**
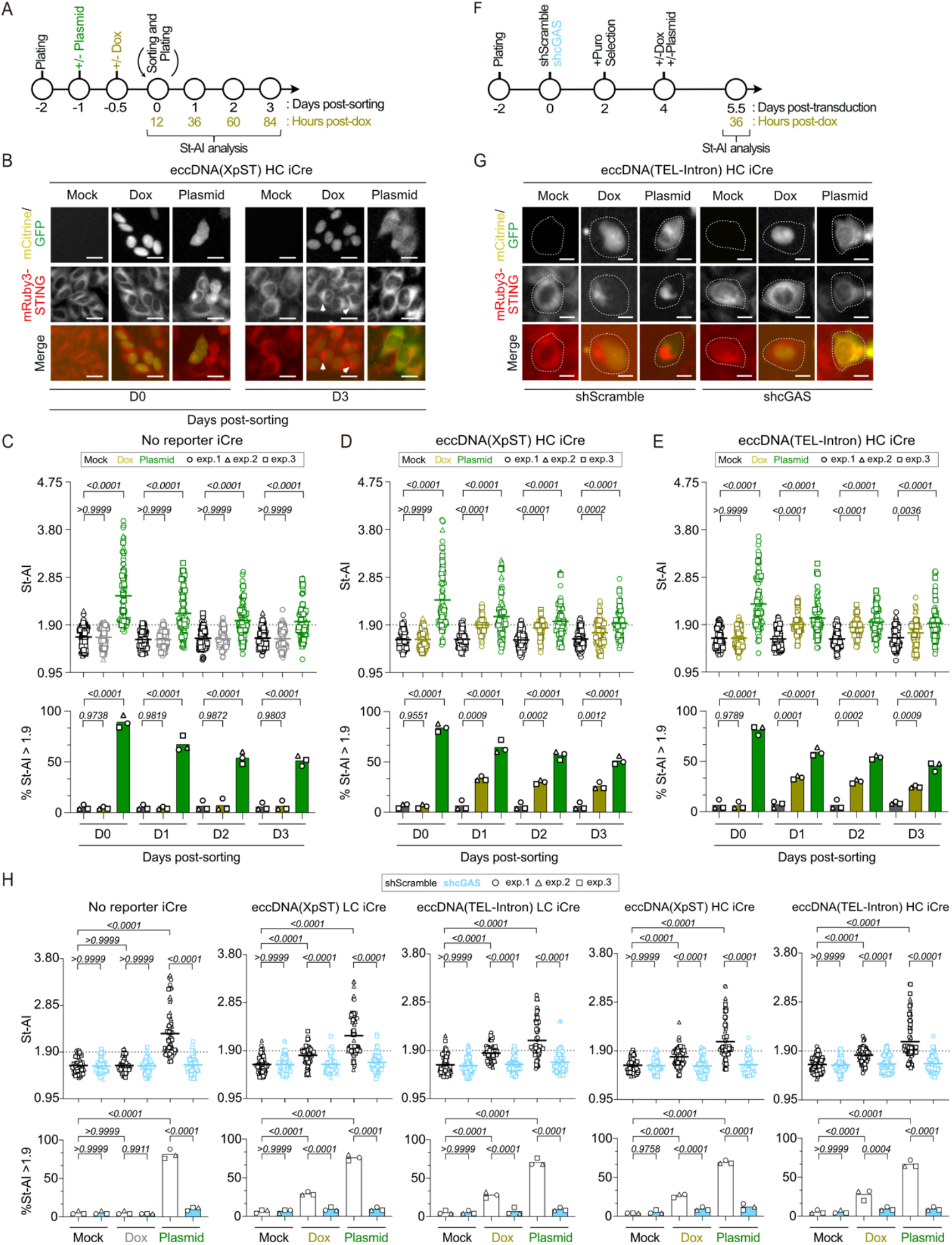
Chromosome-derived eccDNA induces modest cGAS-dependent STING accumulation. **(A)** Experimental workflow for St-AI analysis following sorting of mCitrine-positive eccDNA reporter cells or GFP-positive plasmid-transfected cells. **(B)** Representative images of mCitrine (mock and dox), GFP (plasmid), and mRuby3-STING in HeLa eccDNA(XpST) HC iCre mRuby3-STING cells following mock treatment, dox treatment, or plasmid transfection at days 0 and 3 post-sorting. Arrowheads indicate cells with modestly elevated St-AI. Scale bars: 25 μm. **(C-E)** Top: St-AI analysis of HeLa no reporter iCre (C), HeLa eccDNA(XpST) HC iCre (D), and HeLa eccDNA(TEL-Intron) HC iCre (E) mRuby3-STING cells following the workflow shown in A. Bottom: the corresponding percentages of cells with St-AI > 1.9 are shown as bar graphs. **(F)** Experimental workflow for cGAS knockdown and St-AI analysis of HeLa eccDNA reporter iCre cells. **(G)** Representative images of mCitrine (mock and dox), GFP (plasmid), and mRuby3-STING in shScramble- and shcGAS-transduced HeLa eccDNA(TEL-Intron) HC iCre mRuby3-STING cells following mock treatment, 36 h dox treatment, or 36 h GFP-plasmid transfection. Scale bars: 25 μm. **(H)** Top: St-AI analysis of indicated mRuby3-STING cells following the workflow shown in F. Bottom: the corresponding percentages of cells with St-AI > 1.9 are shown as bar graphs. **Data information: (C-E, H)** Bold lines represent the mean of three independent biological replicates. (**C-E**, n = 100 cells in total, 50 cells in replicate 1 and 25 cells each in replicates 2 and 3; **H**, n = 75 cells in total, 25 cells/biological replicate). Statistical significance was determined by the Kruskal-Wallis test followed by Dunn’s multiple-comparison test (C-E, H: Top) and by ordinary one-way ANOVA followed by Dunnett’s multiple-comparison test (C-E: Bottom) or Šídák’s multiple-comparison test (H: Bottom). P values are indicated for each comparison.

### STING accumulation induced by chromosome-derived eccDNA is uncoupled from its activation

To determine whether the modest St-AI elevation observed in eccDNA reporter cells reflects STING activation, cells were fixed on each day after sorting and immunostained for phospho-STING (Ser366) (pSTING-S366), a marker of STING activation (Tanaka & Chen, 2012; Liu et al., 2015). pSTING-S366 signal intensity increased together with St-AI at the single-cell level in both plasmid-transfected and Compound 3-treated cells (Fig. S7A-E), indicating that an increase in St-AI is accompanied by STING phosphorylation under these conditions. In contrast, none of the eccDNA reporter subclones showed a significant increase in pSTING-S366 signal on days 1, 2, or 3 after sorting (Fig. 6A-D, and S7F, G). To relate STING accumulation to phosphorylation at the single-cell level, cells were classified into four quadrants according to St-AI (threshold, 1.9) and pSTING-S366 intensity (threshold, 2.0) (Fig. 6E, and S7H). At day 0, plasmid-transfected cells were predominantly St-AI-high and pSTING-high, whereas mock- and dox-treated cells were almost exclusively St-AI-low and pSTING-low. At day 1, notably, a St-AI-high and pSTING-low population was prominent in dox-treated eccDNA reporter cells (27.3-31.3%), but rare in mock-treated (1.3-2.0%) and plasmid-transfected (3.3-5.3%) cells (Fig. 6E). Consistent with this, RT-qPCR of downstream chemokines and cytokines, including *CCL5*, *CXCL10*, and *IL-6*, in RNA from the same cells showed no upregulation of these genes at days 1 or 3 post-sorting in any dox-treated reporter subclone (Fig. 6F). Together, these results indicate that STING signaling downstream of its accumulation is not engaged in cells carrying chromosome-derived eccDNA.

**Figure 6.**
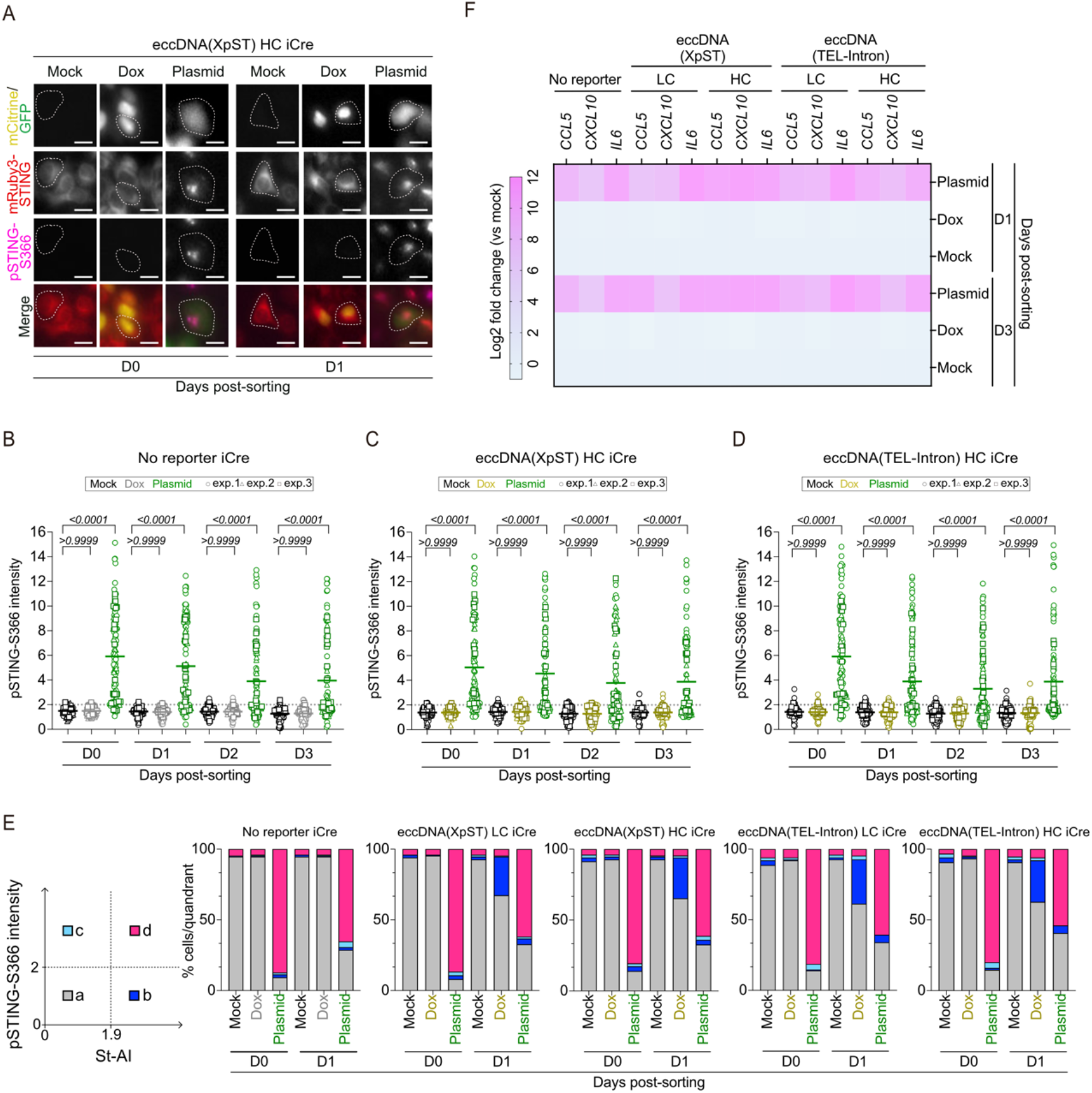
STING accumulation induced by chromosome-derived eccDNA is uncoupled from its activation. **(A)** Representative images of mCitrine (mock, dox), GFP (plasmid), mRuby3-STING, and pSTING-S366 immunofluorescence (IF) in HeLa eccDNA(XpST) HC iCre mRuby3-STING cells on days 0 and 1 post-sorting following mock treatment, dox treatment, or plasmid transfection, as shown in Fig. 5A. White dotted lines indicate cell boundaries. Scale bars: 25 μm. **(B-D)** pSTING-S366 intensity in HeLa no reporter iCre (B), HeLa eccDNA(XpST) HC iCre (C), and HeLa eccDNA(TEL-Intron) HC iCre (D) mRuby3-STING cells following the workflow shown in Fig. 5A. **(E)** Percentages of the indicated HeLa eccDNA iCre cells in each quadrant (a-d) defined by St-AI (threshold, 1.9) and pSTING-S366 (threshold, 2.0), as illustrated in the schematic, on days 0 and 1 post-sorting. The corresponding plots of individual cells are shown in Fig. S7H. **(F)** Heatmap of cytokine and chemokine mRNA expression levels in the indicated HeLa eccDNA reporter iCre cells on days 1 and 3 post-sorting. Expression was normalized to *GAPDH*, and the color scale represents log2 fold change relative to mock-treated cells. **Data information: (B-D)** Bold lines represent the mean of three independent biological replicates (n = 100 cells in total, 50 cells in replicate 1 and 25 cells each in replicates 2 and 3). **(F)** Heatmap values represent the mean of three independent biological replicates. Statistical significance in (B-D) was determined by the Kruskal-Wallis test followed by Dunn’s multiple-comparison test. P values are indicated for each comparison.

### TREX1 does not limit eccDNA-induced STING accumulation

To examine whether the cytosolic exonuclease TREX1 limits eccDNA-induced STING accumulation, we transduced the eccDNA iCre reporter subclones with a lentivirus encoding shTREX1-A before dox treatment (Fig. 7A and S8A). shTREX1-A transduction did not affect St-AI in dox-treated reporter cells (Fig. 7B-E and S8B, C). As a control, treatment with the MPS1 kinase inhibitor reversine induced a modest St-AI elevation comparable to that after 36 hours of dox treatment, and this elevation was further increased by shTREX1-A, as shown by the percentage of St-AI-high cells in all cell lines tested (Fig. 7C-E and S8B, C). This result is consistent with previous findings that TREX1 limits STING activation in the context of drug-induced genotoxic stress (Sato & Hayashi, 2024) and confirms that TREX1 knockdown was functional in these cell lines. Together, these results suggest that TREX1 is dispensable for limiting STING accumulation upon chromosome-derived eccDNA leakage into the cytosol.

**Figure 7.**
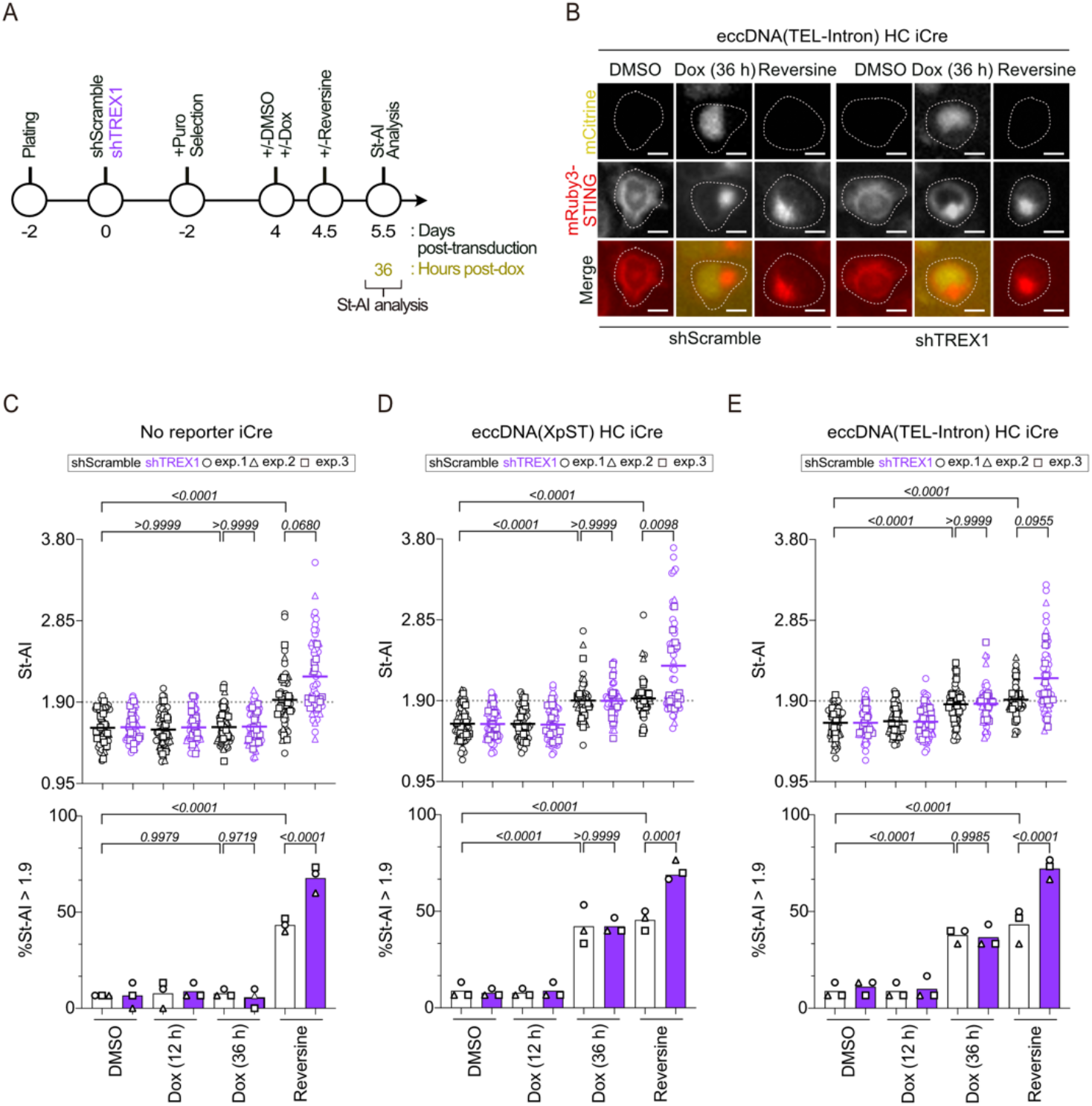
TREX1 does not limit eccDNA-induced STING accumulation. **(A)** Experimental workflow for TREX1 knockdown and St-AI analysis. **(B)** Representative images of mCitrine and mRuby3-STING in shScramble- and shTREX1-transduced HeLa eccDNA(TEL-Intron) HC iCre mRuby3-STING cells following DMSO treatment, 36 h dox treatment, or 24 h reversine treatment. Scale bars: 25 μm. **(C-E)** Top: St-AI analysis of HeLa no reporter iCre (C), HeLa eccDNA(XpST) HC iCre (D), and HeLa eccDNA(TEL-Intron) HC iCre (E) mRuby3-STING cells following DMSO treatment, 12 h and 36 h dox treatment, or 24 h reversine treatment in shScramble- or shTREX1-transduced cells. Bottom: the corresponding percentages of cells with St-AI > 1.9 are shown as bar graphs. For the 12 h dox condition, only mCitrine-positive cells were analyzed, whereas all cells were analyzed for the 36 h dox and reversine conditions. **Data information: (C-E)** Bold lines and bars represent the mean of three independent biological replicates (n = 60 cells in total, 30 cells in replicate 1 and 15 cells each in replicates 2 and 3). Statistical significance was determined by the Kruskal-Wallis test followed by Dunn’s multiple-comparison test (C-E: Top) and by ordinary one-way ANOVA followed by Šídák’s multiple-comparison test (C-E: Bottom). P values are indicated for each comparison.

## Discussion

In this study, we developed a Cre-loxP-dependent eccDNA reporter system that enables the inducible generation of chromosome-derived eccDNA without exogenous genotoxic stress, while simultaneously permitting single-cell-level tracking of eccDNA formation through a fluorescent reporter. Using this system, we showed that chromosome-derived eccDNA leaks into the cytosol and induces a modest, cGAS-dependent accumulation of STING. However, this accumulation was not accompanied by STING phosphorylation at Ser366, nor by upregulation of downstream cytokines and chemokines, in any of the reporter configurations tested. Chromosome-derived eccDNA thus differs from micronuclei, whose chromatin DNA fails to activate cGAS (Sato & Hayashi, 2024; Takaki et al., 2024; Smarduch et al., 2025; Lebrec et al., 2026): eccDNA engages cGAS productively, yet the resulting signal is insufficient to complete the STING activation cycle. These findings suggest that chromosome-derived eccDNA is a weak rather than an inert activator of the cGAS-STING pathway, and that pathway output is restricted at more than one step.

### Possible mechanisms restricting cGAS-STING pathway activation by chromosome-derived eccDNA

Several mechanisms may account for the lack of STING phosphorylation and downstream gene induction by chromosome-derived eccDNA. Importantly, the restriction is not absolute at the level of cGAS: eccDNA induction caused a reproducible, cGAS-dependent elevation of St-AI (Fig. 5H), indicating that cytosolic eccDNA engages cGAS productively enough to drive STING trafficking and accumulation. This distinguishes eccDNA from MN, where cGAS localizes to ruptured structures without detectable cGAMP production or downstream phosphorylation (Sato & Hayashi, 2024; Takaki et al., 2024; Smarduch et al., 2025; Lebrec et al., 2026). Chromosome-derived eccDNA therefore behaves as a partial agonist of the pathway, and the mechanisms discussed below are best understood as quantitative attenuators acting at cGAS, together with a distinct restriction operating downstream of STING accumulation.

Because eccDNA in our system originates from a genome-integrated locus, it is expected to retain nucleosomes upon circularization, and chromatinization is most likely the source of attenuation at cGAS. Structural and cryo-electron microscopy studies have shown that interactions between cGAS and histone H2A-H2B dimers occlude the principal DNA-binding surface of cGAS and sterically inhibit its oligomerization into a catalytically active complex (Boyer et al., 2020; Cao et al., 2020; Kujirai et al., 2020; Michalski et al., 2020; Pathare et al., 2020; Zhao et al., 2020). In addition, barrier-to-autointegration factor 1 (BAF) dynamically outcompetes cGAS for binding to exposed genomic DNA and is required to restrain cGAS upon acute loss of nuclear membrane integrity (Guey et al., 2020); whether BAF acts on cytosolic eccDNA is unknown.

We also note that St-AI did not scale with the genomic reporter copy number: HC subclones, which carry approximately ten-fold more reporter copies than LC subclones, did not show a proportionally greater St-AI response (Fig. 5D, E, 6E, and S6A, B). This is not attributable to saturation for the readout, since the fraction of St-AI-high cells continued to increase with plasmid dose well above the levels reached after eccDNA induction (Fig. S3B-E). The response therefore appears to be limited by a step that is not relieved by supplying more eccDNA, although our cytosolic measurements do not resolve whether the amount of eccDNA reaching the cytosol scales with genomic copy number. That eccDNA nonetheless elicits a measurable cGAS-dependent response suggests that nucleosome-mediated inhibition is not absolute. One possibility is that transcription of the circularized reporter, which is required for mCitrine expression and therefore occurs in every eccDNA-positive cell scored, transiently evicts nucleosomes and exposes DNA to cGAS.

TREX1, a cytosolic 3ʹ→5ʹ exonuclease known to limit the innate immune response to mislocalized self-DNA (Stetson et al., 2008), did not alter the St-AI elevation induced by eccDNA, although the same knockdown enhanced the response to reversine (Fig. 7 and S8). Its lack of effect suggests that cytosolic eccDNA remains largely topologically closed, rather than being nicked or linearized after entering the cytosol. This is of interest because purified eccDNA is a potent STING-dependent immunostimulant in a manner dependent on its circularity (Wang et al., 2021). Our system differs from that study in several respects –– the eccDNA here is chromosome-derived and transcriptionally active, is generated endogenously rather than transfected, and is assayed in an epithelial rather than a myeloid cell –– so the comparison cannot isolate a single determinant. What it indicates is that circularity alone is not sufficient for potent activation. Circularity may contribute positively in our system by conferring resistance to exonucleolytic degradation, while the chromatinized state limits cGAS engagement; the modest response we observe may reflect the balance of the two. Taken together, these observations constrain where the restriction lies. The stimulatory ligand persists in the cytosol in its circular form, and cGAS is catalytically engaged with it. The absence of STING phosphorylation therefore cannot be explained by cGAS-level restriction alone, and points to a second checkpoint downstream of STING accumulation.

### eccDNA uncouples STING accumulation from its activation

In the current mechanistic framework, STING signals only after arriving at the Golgi, where TBK1 recruitment and STING phosphorylation occur (Jeltema et al., 2023). Golgi arrival is necessary but not sufficient: STING that has translocated normally still fails to signal when palmitoylation or the Golgi lipid environment is disrupted (Mukai et al., 2016; Takahashi et al., 2021). To our knowledge, however, such uncoupling has not previously been described for a physiological DNA ligand; prior demonstrations have relied on pharmacological or mutational blockade of a downstream step. Our data show that chromosome-derived eccDNA induces a cGAS-dependent accumulation of STING that is not accompanied by phosphorylation at Ser366 or by immune gene induction, indicating that STING accumulation alone is insufficient for its activation. The comparison with transfected plasmid DNA supports this interpretation. At the threshold plasmid dose, 44– 48% of cells were St-AI-high (St-AI > 1.9) (Fig. 3G), comparable to the 25–42% observed after eccDNA induction (Fig. 5D, E, 7D, E, S6A, B, and S8B, C). However, *CCL5* was induced only by plasmid transfection (Fig. 3C, 6F), whereas *CCL5*, *CXCL10*, and *IL-6* were not upregulated in eccDNA reporter cells (Fig. 6F). These observations suggest that the restriction in the eccDNA condition operates not only at the level of cGAS activation, but also at a subsequent step in the STING activation cycle.

One possibility is that the restriction acts on STING itself at the Golgi. As described above, arrival at the Golgi is not merely positional: STING activation requires palmitoylation at Cys88/91, and pharmacological or genetic blockade of this modification abolishes the type I interferon response even though STING has translocated normally (Mukai et al., 2016). Consistent with this, a cell-free reconstitution showed that TBK1-mediated STING phosphorylation depends on ER-to-Golgi traffic, on palmitoylation, and on the sphingomyelin- and cholesterol-rich lipid environment of the trans-Golgi network (Takahashi et al., 2021). Accumulation of STING at the Golgi therefore need not imply that it has entered a signaling-competent state. Whether eccDNA-induced STING accumulation is accompanied by palmitoylation, and whether the accumulated pool resides in the appropriate trans-Golgi subcompartment, remain to be determined.

A second, non-exclusive possibility concerns the magnitude of STING mobilization rather than a discrete downstream block. IFI16 is required for cGAMP-induced STING activation and interacts with STING to promote both its phosphorylation and its translocation from the ER; in IFI16-deficient keratinocytes, DNA-induced STING relocalization is reduced (Almine et al., 2017). IFI16 therefore appears to set the extent of STING mobilization rather than to act as a simple switch. If eccDNA-associated cGAS complexes engage IFI16 only weakly, the resulting STING accumulation may be submaximal: sufficient to exceed the St-AI threshold in a substantial fraction of cells, yet insufficient to reach the local STING density required for productive signaling. This connects directly to the structural requirement for phosphorylation, since cryo-electron microscopy of the STING-TBK1 complex showed that Ser366 in the STING C-terminal tail cannot reach the kinase active site of the bound TBK1 dimer, so STING phosphorylation requires cGAMP-induced higher-order oligomerization of both proteins (C. Zhang et al., 2019). Whatever the precise mechanisms are, the lack of STING activation was not affected by the two inserted sequences tested: the presence of telomeric repeats had no discernible effect on the cGAS-STING response to chromosome-derived eccDNA.

### Perspectives

The Cre-loxP-based eccDNA reporter system established here provides a well-controlled platform for dissecting the innate immune consequences of chromosome-derived circular DNA independently of the genotoxic and mitochondrial confounders inherent to stress-based eccDNA induction strategies. The principle underlying this approach, decoupling the generation of a specific self-DNA species from mitochondrial perturbation, may be applicable beyond eccDNA. Other endogenous nucleic acid species proposed as innate immune activators have been similarly studied in contexts where mitochondrial confounders cannot be excluded.

For example, R-loop-derived cytoplasmic RNA-DNA hybrids have been shown to activate cGAS and TLR3 in SETX- and BRCA1-deficient cells (Crossley et al., 2023). Yet SETX depletion alone is sufficient to induce R-loop accumulation in the regulatory region of mitochondrial DNA and to elevate intracellular ROS, and comparable mitochondrial defects accompany BRCA2 deficiency (Renaudin et al., 2021).

Similarly, LINE-1 reverse transcription intermediates have been implicated as cGAS activators, both in senescent cells (De Cecco et al., 2019) and following pharmacological L1 de-repression (Baldwin et al., 2024). Interpretation in both settings is complicated by routes to the same sensor that bypass L1. In the senescence model, the triple intervention used to reconstitute L1 activation includes TREX1 knockdown, which independently relieves cGAS-STING restriction, and senescent cells release mtDNA into the cytosol through BAX/BAK macropores to activate cGAS-STING (Victorelli et al., 2023). Where L1 dependence is inferred from derepression of L1 by decitabine and its reversal by nucleoside reverse transcriptase inhibitors, the inference is likewise not clean: decitabine raises ROS and dissipates mitochondrial membrane potential, an effect requiring incorporation into DNA and therefore inseparable from demethylation (Shin et al., 2012; Fandy et al., 2014). Whether this releases mtDNA is untested, but mitochondrial permeabilization activates cGAS-STING (McArthur et al., 2018; Victorelli et al., 2023), and reversal by reverse transcriptase inhibitors cannot distinguish the two sources, since their triphosphates also inhibit mitochondrial DNA polymerase and may deplete mtDNA (Lewis et al., 2001). Rigorous re-evaluation of the intrinsic immunogenicity of these nucleus-derived nucleic acid species in living cells, using approaches that decouple their induction from mitochondrial perturbation, will be essential for advancing our understanding of self-nucleic acid sensing in genome instability, cellular senescence, and autoimmune disease.

## Limitations

Several limitations of the present study warrant consideration. First, our experiments were conducted primarily in HeLa1.2.11 cells. The extent to which our findings generalize to other cell types remains to be determined. Second, the Cre-loxP system generates eccDNA from a defined genomic sequence without exogenous genotoxic stress, although Cre-mediated recombination itself involves transient DNA strand breakage and rejoining. While this design is well suited to excluding mitochondrial confounders, naturally arising eccDNA in cancer cells, which forms through genotoxic stress, may carry distinct histone modifications or higher-order chromatin structures that differ from those of reporter-derived eccDNA. Third, although the lack of a proportional increase in STING accumulation from LC to HC subclones argues against copy number as the primary determinant of the failure to activate STING, the possibility that the substantially higher eccDNA amplicon copy numbers could overcome chromatin-mediated suppression and breach the threshold for full STING activation cannot be excluded.

## Materials and Methods

### Cell culture

Human cervical carcinoma HeLa1.2.11 cells (a gift from Titia de Lange), human colorectal carcinoma HCT116 cells (American Type Culture Collection, ATCC), and human fibrosarcoma HT1080 cells (ATCC), were maintained in Dulbecco’s Modified Eagle Medium (DMEM, 05919; Shimadzu Diagnostics) supplemented with 10% fetal bovine serum (FBS, S1810-500; BioWest), 0.165% NaHCO_3_, 2 mM L-glutamine, 5 μg/mL Plasmocin (InvivoGen), and 100 U/mL penicillin/streptomycin. Cells were incubated at 37°C in a humidified atmosphere with 5% CO_2_. For passaging, the medium was first removed, and cells were rinsed with 1× PBS. A solution containing 0.1 mM EDTA and 0.1% trypsin was added to the dish and incubated at 37°C for 5 minutes. After detachment, cells were resuspended in fresh complete medium. Doses and durations other than 1 µM Compound 3 (diABZI STING agonist-1, S8796; Selleck) and 1 µg/mL doxycycline (D9891; Sigma-Aldrich) are indicated in the figure legends. For MPS1 inhibition, cells were treated with 0.5 μM reversine (10004412; Cayman Chemical) for 24 hours.

### Plasmid construction

All plasmid clonings were first modeled in SnapGene software (version 5.0.8) and then performed using conventional restriction digestion and/or PCR, followed by NEBuilder HiFi DNA assembly (NEB). Briefly, the mRuby3-hSTING coding sequence from pMX-mRuby3-hSTING (Sato & Hayashi, 2024) was cloned into a third-generation lentiviral vector to obtain pLenti-mRuby3-hSTING (pMTH1770). The pCWtetV16 vector was generated from the pCW-Cas9, originally obtained from the E. Lander and D. Sabatini laboratories (addgene #50661), by introducing the rtTA-V16 variant (Zhou et al., 2006; Das et al., 2016). The Cre gene from pMTH225/pGFAP-Cre (a gift from A. Messing) was then introduced into this vector to obtain pCWtetV16-Cre (pMTH1826). A Sleeping Beauty donor vector, pSBCm (pMTH1889), was generated by integrating artificially synthesized 5ʹ-ITR and 3ʹ-ITR sequences (Ivics et al., 1997) into the pCm vector (pMTH1747). All eccDNA reporter donor cassettes were first constructed on the pBSII-NotI vector (pMTH295) and re-cloned into the NotI site of pSBCm. The Fusion Visualization (FuVis) reporter cassette, pMTH397 (Kagaya et al., 2020), was used as a template to generate pBS-P-mCitC-N-P-neoR (pMTH1824) and pBS-P-mCitC-XpSubTel691-N-P-neoR (pMTH1864). These constructs were then used to obtain the corresponding SB donor vectors, pMTH1896 and pMTH1899, respectively. A plasmid encoding 114 tandem repeats of the TTAGGG telomere sequence interrupted by 13 bp non-telomeric spacer sequences (pMTH1808/pSV40_TelNN; a gift from Y. Doksani) (Huda et al., 2023) was used to integrate telomere repeats into the eccDNA(-) cassette. To clone pSBCm-P-mCitC-TelNN691-N-P-neoR (pMTH1898), Esp3I-ClaI sites were introduced into the NheI site upstream of the EF1α promoter on pMTH1824 to obtain pMTH1831, into which the Esp3I-ClaI fragment of pMTH1808 was introduced to obtain pBS-P-mCitC-TelNN691-N-P-neoR (pMTH1853); pMTH1853 was then re-cloned into the NotI site of pSBCm to obtain pMTH1898. To clone pSBCm-P-mCitC-N-TelNN691-P-neoR (pMTH1996), SpeI-BamHI-ClaI-EcoRI sites were introduced into the EcoRI site of pMTH1824 to obtain pMTH1930. In parallel, NheI-SpeI-BspEI-SpeI-NheI sites were introduced into the ClaI site of pMTH1853 to obtain pMTH1942. Subsequently, a 730 bp NheI fragment of pMTH1942 was cloned into the NheI site of pMTH1930 to obtain pBS-P-mCitC-N-TelNN691-P-neoR (pMTH1987); pMTH1987 was then re-cloned into the NotI site of pSBCm to obtain pMTH1996. pMTH1896, pMTH1899, pMTH1898, and pMTH1996 were used to establish the HeLa eccDNA(-), eccDNA(XpST), eccDNA(TEL), and eccDNA(TEL-Intron) reporter cells, respectively. Plasmids used in this study are listed in Table 1 and the full-length sequences of the plasmids made in this study are available at a public data-sharing server (DOI:10.6084/m9.figshare.34745058).

### Sleeping Beauty-mediated reporter integration

Stable integration of the eccDNA reporter cassette was performed using the Sleeping Beauty (SB) transposon system as previously described (Luqman-Fatah et al., 2023) with minor modifications. Briefly, 2 × 10^5^ HeLa1.2.11, HT1080, and HCT116 cells were seeded in 6-well plates and co-transfected with a plasmid encoding the SB transposase (pMTH1633; 50 ng) and one of the eccDNA reporter donor plasmids (500 ng) using FuGENE HD transfection reagent (Promega) in Opti-MEM according to the manufacturer’s instructions. After 24 hours, transfected cells were subjected to G418 selection (500 μg/mL for HeLa1.2.11 and HT1080 cells and 700 μg/mL for HCT116 cells) for 14 days, followed by no selection for 7 days and subsequent G418 selection at the same concentrations for 7 days to obtain stable integrants. Integration of reporter cassettes was validated by genomic PCR. Stable cell populations were subsequently used for single-cell cloning and downstream analyses.

### Single-cell cloning of eccDNA reporter cells

To establish clonal eccDNA reporter cell lines, stable HeLa1.2.11 eccDNA reporter cells were collected by trypsinization, resuspended in culture medium, and diluted in 1× PBS containing 0.1 mM EDTA to obtain a sparse suspension of individual cells. Single cells were manually isolated using a micropipette under a microscope with 10× objective and transferred into individual wells of a 96-well plate containing culture medium. Cells were cultured for approximately 2 weeks, and wells containing colonies derived from single cells were identified by microscopic observation. When colonies reached an appropriate size, cells were expanded sequentially into larger culture vessels and cryopreserved. Candidate clones were further evaluated based on mCitrine induction upon Cre transduction and reporter copy number.

### Validation of reporter integration by genomic PCR

To validate integration of the eccDNA reporter cassette, genomic DNA was extracted from target cell lines using the Wizard® Genomic DNA Purification Kit (Promega) according to the manufacturer’s instructions. Genomic PCR was performed using Tks Gflex DNA Polymerase (Takara) and the primer sets listed in Table 2. PCR products were analyzed by agarose gel electrophoresis to confirm integration of the reporter cassette.

### Quantification of reporter copy number

Genomic DNA was extracted as described above, and the copy number of the integrated reporter was determined by qPCR using primers specific for mCitrine and the AAVS1 locus (Table 2). Reporter copy number was expressed relative to the endogenous AAVS1 locus (AAVS1 = 1), calculated from the mCitrine/AAVS1 qPCR ratio normalized to that of the reference plasmid pMTH864, which contains one copy of each sequence. Because the AAVS1 copy number in HeLa cells was not determined, values can be converted to absolute copy numbers per genome by multiplying by the AAVS1 copy number. Subclones carrying relative copy numbers approximately 4–13 and 70–150 were designated low-copy (LC) and high-copy (HC), respectively.

### Rolling circle amplification

Rolling Circle Amplification (RCA) was conducted as previously described (Dean et al., 2001) with minor modifications. Briefly, for a 20 μL RCA reaction, the following components were mixed: 2 μL of 10x Phi29 DNA polymerase reaction buffer (Takara), 4 μL of primer mixture containing 1 μL each of four primers (100 μM; MTH514, MTH697, MTH1283, and MTH3048; see Table 2), 4 μL of 10 mM dNTPs (Takara), and 70 ng of DNA template. Nuclease-free water was added to bring the total volume to 18.6 μL. The reaction mixture (without polymerase) was heated at 95°C for 5 minutes to denature circular DNA. After 1 minute of cooling at room temperature, 1 μL of Phi29 DNA polymerase (Takara), 0.2 μL of 100 mM DTT, and 0.2 μL of BSA were added. The reaction mixture was incubated at 30°C for 16 hours, followed by 65°C for 30 minutes to inactivate the enzyme.

### Virus transduction

Lentiviral particles were generated as previously described (Sato & Hayashi, 2024) with minor modifications. Briefly, 1.6 μg of transfer plasmid was co-transfected into Lenti-X 293T cells (632180, Takara Bio) with 0.8 μg of the packaging plasmid psPAX2 (#12260; Addgene, a gift from Didier Trono) (pMTH688) and 0.8 μg of the envelope plasmid pCMV-VSV-G (#8454; Addgene, a gift from Robert A. Weinberg) (pMTH689) using 9.6 μL of 1 mg/mL polyethylenimine (PEI) in a 6-well plate containing antibiotic-free medium. The medium was replaced with fresh medium containing antibiotics 24 hours post-transfection. Lentivirus-containing supernatants were collected on days 2 and 3 post-transfection and filtered through a 0.45-μm PES syringe filter (TELS25045; Technolabs Inc.). For lentiviral transduction, the culture medium of target cells was replaced with virus-containing medium supplemented with 8 μg/mL polybrene. For constitutive Cre expression, cells were transduced with a lentivirus encoding Cre recombinase produced from pLenti-Cre (pMTH1278) or pLenti-bla-p2a-emiRFP703 (pMTH1050) as a Lentivirus control (Fig. S4A). To generate mRuby3-STING reporter-expressing cells, eccDNA reporter cells were transduced with a lentivirus produced from pLenti-mRuby3-hSTING (pMTH1770) and enriched by FACS as described in ‘Flow cytometry’. For shRNA-mediated knockdown of TREX1 and cGAS, cells were transduced with lentiviruses produced from pLKO.1-shTREX1-A (pMTH1657), pLKO.1-shcGAS-1 (pMTH1502), or pLKO.1-shScramble (pMTH127). Transduced cells were selected with 1 μg/mL puromycin (ChemCruz) starting 2 days post-transduction and maintained under selection for 2 days. For the shcGAS-1 and shTREX1-A experiments, lentiviral transduction was performed 4 days before dox treatment or plasmid transfection, and reversine was added for 24 hours starting 4.5 days after transduction. The following shRNA sequences were used (5′ to 3′): shScramble, CCTAAGGTTAAGTCGCCCTCG; shTREX1-A, AACACGGCCCAAGGAAGAGCT; and shcGAS-1, TTAGTTTTAAACAATCTTTCCT. To generate dox-inducible Cre (iCre) cells, HeLa cells and HeLa eccDNA reporter subclones expressing mRuby3-hSTING were transduced with a lentivirus encoding pCWtetV16-Cre (pMTH1826) and selected with 10 μg/mL blasticidin starting 2 days post-transduction for at least 3 days.

### Plasmid transfection for STING assay

Cells were seeded in 6-well plates at a density of 2 × 10⁵ cells per well. On the following day, cells were transfected with pMAX-TurboGFP (pMTH380) plasmid DNA using FuGENE HD transfection reagent (Promega). Briefly, 1 μg plasmid DNA was mixed with 2 μL FuGENE HD in 100 μL Opti-MEM (Thermo Fisher Scientific), incubated for 30 minutes at room temperature, and added to the cells in 1.9 mL medium. Following transfection, cells were incubated for indicated duration at 37°C under standard culture conditions before subsequent experiments. For plasmid titration experiments, serial two-fold dilutions ranging from 2^0^ to 2^-11^ μg plasmid DNA per 2 mL medium were used, with FuGENE HD added at a constant ratio of 2 μL per μg plasmid DNA.

### Flow cytometry

Flow cytometry was performed as previously described (Sato & Hayashi, 2024) with minor modifications. Briefly, cells were harvested by trypsinization, washed with 1× PBS, and resuspended in cold 1× PBS containing 0.1 mM EDTA before filtration through a 5 mL polystyrene round-bottom tube with a cell-strainer cap (352235, Corning). Cells were analyzed using an SH800S Cell Sorter (Sony) equipped with a 100 μm or 130 μm sorting chip. Single cells were gated based on forward scatter width (FSC-W) values prior to analysis and sorting. Fluorescence signals were detected using the following laser and filter combinations: mCitrine (488 nm laser, 525/50 filter), mRuby3 (561 nm laser, 600/60 filter), and GFP (488 nm laser, 525/50 filter). mRuby3-expressing cells were enriched by three sequential rounds of sorting, with cells expanded in culture between each round before the subsequent sort. For St-AI analysis after sorting, mCitrine-positive cells were sorted 12 hours after the addition of dox, with the gate set using mock-treated cells; GFP-positive cells were sorted from plasmid-transfected cultures 24 hours after transfection. Sorted cells were plated without dox and analyzed on days 1, 2, and 3.

### RNA extraction and RT-qPCR

Total RNA was extracted from cells using the RNeasy Mini Kit (QIAGEN). A total of 0.33 μg of RNA was reverse-transcribed in a 25 μL reaction mixture containing 1× AMV RT reaction buffer (00054B, NIPPON GENE), 0.8 mM dNTPs (Toyobo), 5 mM MgCl₂ (Toyobo), 0.125 μM oligo(dT) primer (Eurofins), and 3.6 units AMV reverse transcriptase (30005G, NIPPON GENE). Reverse transcription was performed at 30°C for 10 minutes, 42°C for 30 minutes, and 95°C for 5 minutes. The resulting cDNA was used for qPCR with SYBR™ Green qPCR Master Mix (C19020F4, Thermo Fisher Scientific). qPCR amplification was performed using a StepOnePlus Real-Time PCR System (Applied Biosystems) with an initial denaturation at 95°C for 15 seconds, followed by 40 cycles of 95°C for 15 seconds and 60°C for 60 seconds. Relative gene expression levels were calculated using the quantity means generated by the StepOnePlus software based on standard curve analysis of Ct values. For each sample, average target gene quantities from two technical replicates were normalized to that of *GAPDH*, and the resulting values were further normalized to the corresponding mock-treated control. For the heatmaps of cytokine and chemokine genes, the mean log2 fold changes from three biological replicates were plotted in GraphPad Prism (version 11.0.2). The primers used for RT-qPCR are listed in Table 2.

### Genomic qPCR

Genomic, cytosolic, and RCA-derived DNA samples were analyzed by qPCR using SYBR Green qPCR Master Mix on a StepOnePlus Real-Time PCR System under the conditions described for RT-qPCR. Quantities were calculated from standard curves generated using serial dilutions of plasmid DNA containing the target sequences. For RCA, mCitrine signals were normalized to neoR and to the corresponding reaction without phi29 DNA polymerase. For cytosolic fractions, signals from 1× 10^6^ cells equivalents of the cytosolic fraction and of whole-cell DNA were normalized to the corresponding mock-treated whole-cell sample.

### Cytosolic fraction and whole-cell DNA isolation

Cytosolic fractions were isolated as previously described (Sato & Hayashi, 2024) with minor modifications. Briefly, cells were treated with dox for 12 or 36 hours, or transfected with the eccDNA donor plasmid (pMTH1899 or pMTH1996) for 24 hours, before collection. A total of 1 × 10^6^ cells were resuspended in 100 μL of digitonin buffer (150 mM NaCl, 50 mM HEPES, pH 7.4, and 25 μg/mL digitonin) and incubated on a rotator for 10 minutes at 4°C. Lysates were centrifuged at 2,000 × g for 10 minutes at 4°C, and the supernatant was transferred to a new tube. The supernatant was further centrifuged at 2,000 × g for 20 minutes at 4°C and transferred to a new tube. This step was repeated three additional times, and the final supernatant was used as the cytosolic fraction. For whole-cell DNA isolation, 1 × 10^6^ cells were harvested under the same experimental conditions, and genomic DNA was extracted using the Wizard® Genomic DNA Purification Kit (Promega) according to the manufacturer’s instructions. Purified DNA was resuspended in 80 μL of DNA Rehydration Solution. Cytosolic and whole-cell DNA samples were subsequently analyzed by genomic qPCR. Primer sequences are listed in Table 2.

### Immunofluorescence

Immunofluorescence was performed as previously described (Sato & Hayashi, 2024) with minor modifications. Briefly, cells were grown on coverslips coated with Alcian Blue 8GX (A5268; Sigma-Aldrich), fixed with 4% paraformaldehyde in 1x PBS for 15 minutes at room temperature, and washed with 1x PBS three times. The fixed cells were permeabilized using 0.2% Triton X-100, 0.02% skim milk (Nacalai), and 0.02% BSA (Sigma-Aldrich) in 1x PBS for 5 minutes at room temperature in the dark. After sequential rinsing with 1x PBS and PBST (0.1% Tween-20, 1x PBS), the cells were incubated with rabbit anti-phospho-STING (Ser366) (19781S; Cell Signaling Technology) antibody diluted in PBST for 45 minutes at room temperature. After three washes with PBST, the cells were incubated with Cy5-labeled goat anti-rabbit IgG (H+L) (PA45004; Cytiva), or Alexa Fluor 488-conjugated goat anti-rabbit (A11034; Invitrogen) (Fig. S7D, E), in PBST for 45 minutes at room temperature in the dark, and then washed with PBST and Milli-Q water. After air drying, coverslips were mounted on glass slides using an in-house antifade mounting medium [4% n-propyl gallate (29303-92, Nacalai), 100 mM Tris-HCl pH 8.7, 90% glycerol (17018-25, Nacalai)]. The primary and secondary antibodies were diluted in PBST as indicated in Table 3.

### Imaging

Live cells expressing GFP, mCitrine, and mRuby3-STING, or fixed cells stained for pSTING-S366 were imaged using a BZ-X710 (Keyence) equipped with a metal halide lamp and a 20×, NA 0.45 objective lens. mCitrine, GFP, Alexa Fluor 488, mRuby3, and Cy5 were detected with the following filter cubes: mCitrine (ex: 500/20 nm, em: 535/30 nm, dichroic: 515LP); GFP and Alexa Fluor 488 (ex: 470/40 nm, em: 525/50 nm, dichroic: 495LP); mRuby3 (ex: 545/25 nm, em: 605/70 nm, dichroic: 565LP); and Cy5 (ex: 620/60 nm, em: 700/75 nm, dichroic: 660LP). Exposure times were set so that no pixels were saturated and were kept constant across conditions within each experiment. Phase contrast images were acquired in parallel to define cell boundaries. Images were acquired from randomly selected fields.

### St-AI and pSTING Analysis

The STING Accumulation Index (St-AI), defined as the ratio of the maximum to the mean mRuby3-STING fluorescence intensity within individual cells, and the mean pSTING immunofluorescence intensity within each cell boundary were quantified using ImageJ Software (v1.53t) as previously described (Sato & Hayashi, 2024). Cell boundaries were defined based on phase contrast image. Thresholds for St-AI (1.9) and pSTING-S366 intensity (2.0) were set at the values that gave the largest difference between mock-treated and plasmid-transfected or Compound 3-treated cells. For the quadrant analysis, cells were classified as St-AI-high or pSTING-high when they exceeded these thresholds.

### Statistical analysis and figure preparation

Statistical analyses were performed using GraphPad Prism (version 11.0.2). Unless otherwise indicated, all experiments were conducted in three independent biological replicates. Individual biological replicates are represented by different symbols in the figures: circles (replicate 1), triangles (replicate 2), and squares (replicate 3). Data from two biological replicates, where indicated, are shown without statistical testing. Single-cell measurements (St-AI and pSTING-S366 intensities) from the three biological replicates were pooled and analyzed by the Kruskal-Wallis test followed by Dunn’s multiple-comparison test. Replicate-level values (%St-AI > 1.9, and genomic and cytosolic-fraction qPCR data) were analyzed by ordinary one-way ANOVA followed by Šídák’s or Dunnett’s multiple-comparisons test, as indicated in the figure legends. The efficiency of shRNA knockdown was analyzed by Welch’s t-test. All tests were two-sided, and a P value (or an adjusted P value for multiple comparisons) < 0.05 was considered statistically significant. The number of cells or biological replicates is given in the figure legends. Graphs were generated in GraphPad Prism (version 11.0.2), and figures were assembled using Affinity Designer V2 (3.3.1) (Serif). Brightness and contrast were adjusted uniformly across entire images.

## Acknowledgments

We thank the Drug Discovery Centre supported by the Innovative Support Alliance for Life Science (iSAL), Kyoto University, for flow cytometry facilities; Ylli Doksani, Zsuzsanna Izsvak, Jan Karlseder, Eric Lander, Titia de Lange, Albee Messing, David Sabatini, Didier Trono, and Robert A. Weinberg for sharing materials; Tomoichiro Miyoshi, Ryota Sato, and Tomohiko Taguchi for sharing materials and unpublished results; and Yumi Hayashi for technical assistance with molecular cloning. We also thank all members of the Hayashi laboratory for their support, suggestions, and discussion.

## Author contributions

H.D. and M.T.H. conceived the study and designed the experiments. H.D. performed most of the experiments and carried out the data analyses. J.C. performed RCA experiments. M.T.H. designed the reporter configuration, contributed to data analysis and interpretation throughout the study, supervised the project, and secured funding. H.D. and M.T.H. wrote the manuscript. All authors reviewed and approved the manuscript.

## Data availability

All data generated or analyzed in this study are available in the article and its Supplementary Information, and from the corresponding author upon reasonable request. DNA sequences of plasmids made in this study are available at a public data-sharing server (DOI: 10.6084/m9.figshare.34745058).

## Ethics approval

This study did not involve human participants or animal models, and therefore did not require ethical approval.

