## Supplementary Figures for "Uncoupling of STING Accumulation and Activation upon Chromosome-Derived Extrachromosomal Circular DNA Formation"

Figure S1: Establishment and validation of eccDNA reporter cells

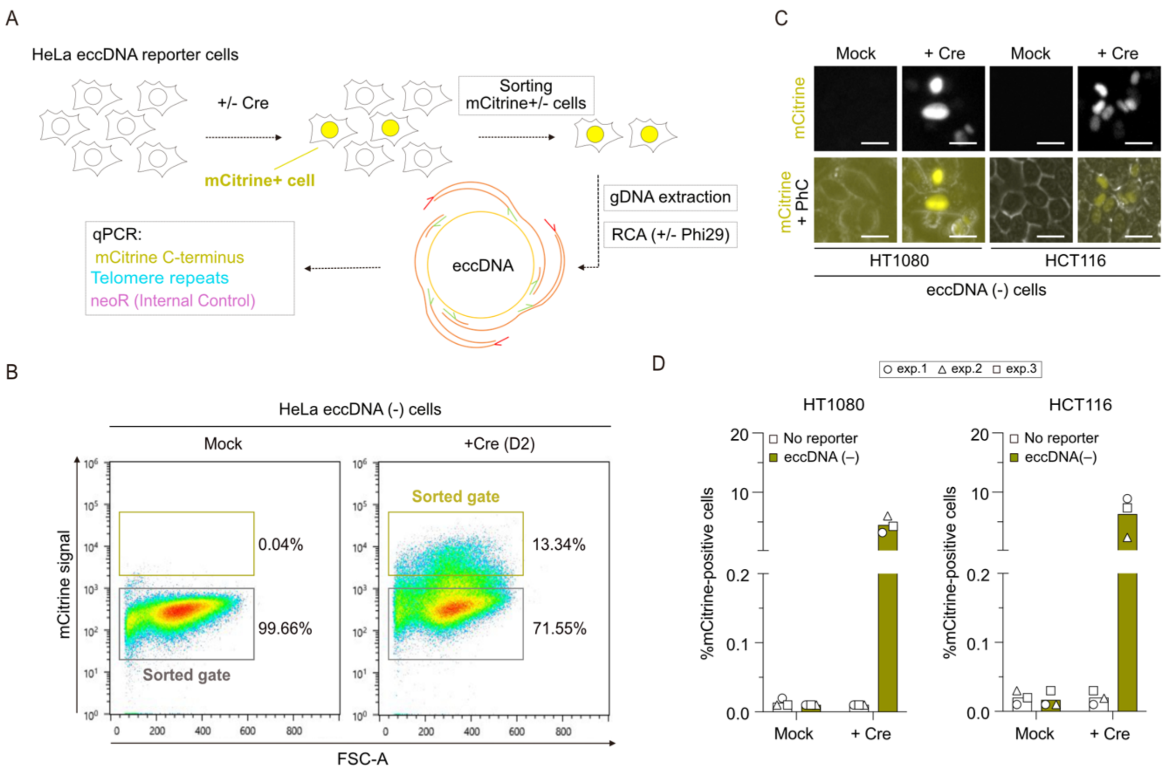


**Figure S1. Establishment and validation of eccDNA reporter cells.**

(A) Experimental workflow for validation of eccDNA formation by rolling circle amplification (RCA). mCitrine-positive cells were sorted 2 days post-Cre transduction and subjected to RCA and qPCR analysis. (B) Representative flow cytometry plots and sorted gates of mCitrine-negative and -positive HeLa eccDNA(-) cells 2 days after mock treatment and Cre transduction, respectively. (C) Representative live-cell images of mCitrine in HT1080 eccDNA(-) and HCT116 eccDNA(-) reporter cells 2 days post-Cre transduction. Mock-treated cells served as negative controls. PhC, phase contrast. Scale bars: 25 µm. (D) Percentages of mCitrine-positive HT1080 eccDNA(-) and HCT116 eccDNA(-) reporter cells 2 days post-Cre transduction. Data information: (D) Bars represent the mean of three independent biological replicates.

Figure S2: Reporter integration and copy-number analysis of eccDNA reporter cells

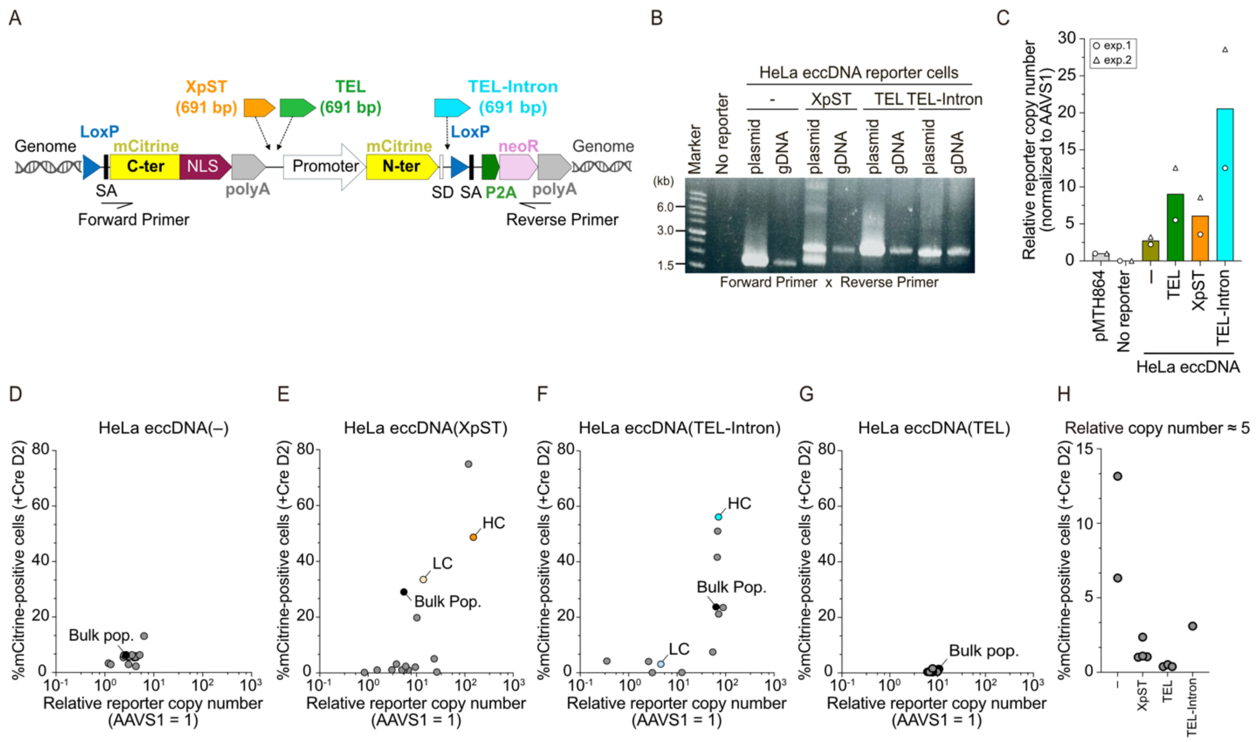


**Figure S2. Reporter integration and copy-number analysis of eccDNA reporter cells.**
(A) Schematic of the Cre-loxP-dependent HeLa eccDNA(XpST), eccDNA(TEL), and eccDNA(TEL-Intron) reporter cell lines. Reporter cassettes were independently integrated into HeLa1.2.11 cells using the Sleeping Beauty transposon system. (B) Representative gel image showing genomic PCR products of reporter cassettes in the indicated HeLa eccDNA reporter cell lines. The corresponding donor plasmids were used as control templates. (C) Relative reporter copy number (AAVS1 = 1) in the indicated HeLa eccDNA reporter cell lines. (D-G) Scatter plots showing the relationship between relative reporter copy number and the percentages of mCitrine-positive cells 2 days post-Cre transduction in single-cell-derived subclones of HeLa eccDNA(-) (D), eccDNA(XpST) (E), eccDNA(TEL-Intron) (F), and eccDNA(TEL) (G) reporter cells. (H) Percentages of mCitrine-positive cells 2 days post-Cre transduction in reporter subclones with a relative copy number of approximately five. Data information: (C) Bars represent the mean of two biological replicates. (D-G) Each symbol represents an individual subclone. LC, low copy; HC, high copy; Bulk Pop., bulk population. (H) n = 2 for eccDNA(-); 4 for eccDNA(XpST); 3 for eccDNA(TEL); and 1 for eccDNA(TEL-Intron).

Figure S3: Development of eccDNA reporter cells equipped with a STING reporter

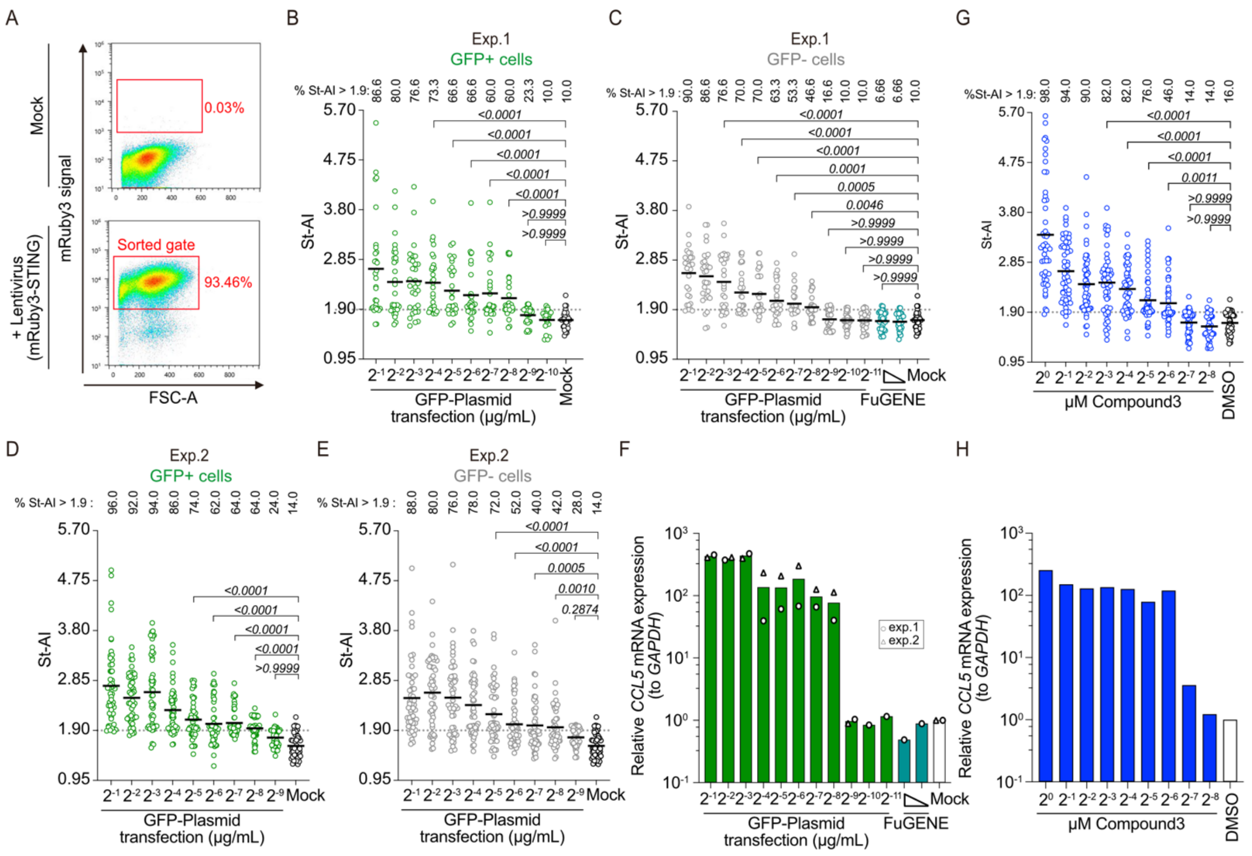


**Figure S3. Development of eccDNA reporter cells equipped with a STING reporter.**
**(A)** Representative flow cytometry plots and a sorted gate of HeLa mRuby3-STING cells following lentiviral transduction. **(B-E)** St-AI analysis in GFP-positive (B, D) and GFP-negative (C, E) cells from experiment 1 (B, C) and 2 (D, E), following transfection with the indicated amounts of a GFP-expressing plasmid or only FuGENE reagent. The same Mock data are shown in B and C, and D and E. **(F)** Relative *CCL5* mRNA expression normalized to *GAPDH* in cells shown in B-E. **(G)** St-AI analysis following treatment with the indicated concentrations of Compound 3 in HeLa mRuby3-STING cells. **(H)** Relative *CCL5* mRNA expression normalized to *GAPDH* in HeLa mRuby3-STING cells in cells shown in (G). **Data information: (B-E, G)** Bold lines represent the mean (B, C, n = 30 cells/treatment; D, E, G, n = 50 cells/treatment). Percentages of cells with St-AI > 1.9 are shown above each plot. **(F)** Bars represent the mean. Data in (F) correspond to the GFP-plasmid transfection experiments shown in (B, C) (experiment 1) and (D, E) (experiment 2). Statistical significance was determined by the Kruskal-Wallis test followed by Dunn's multiple-comparison test. P values are indicated for each comparison.

Figure S4: Establishment and validation of dox-inducible Cre-dependent eccDNA reporter cells

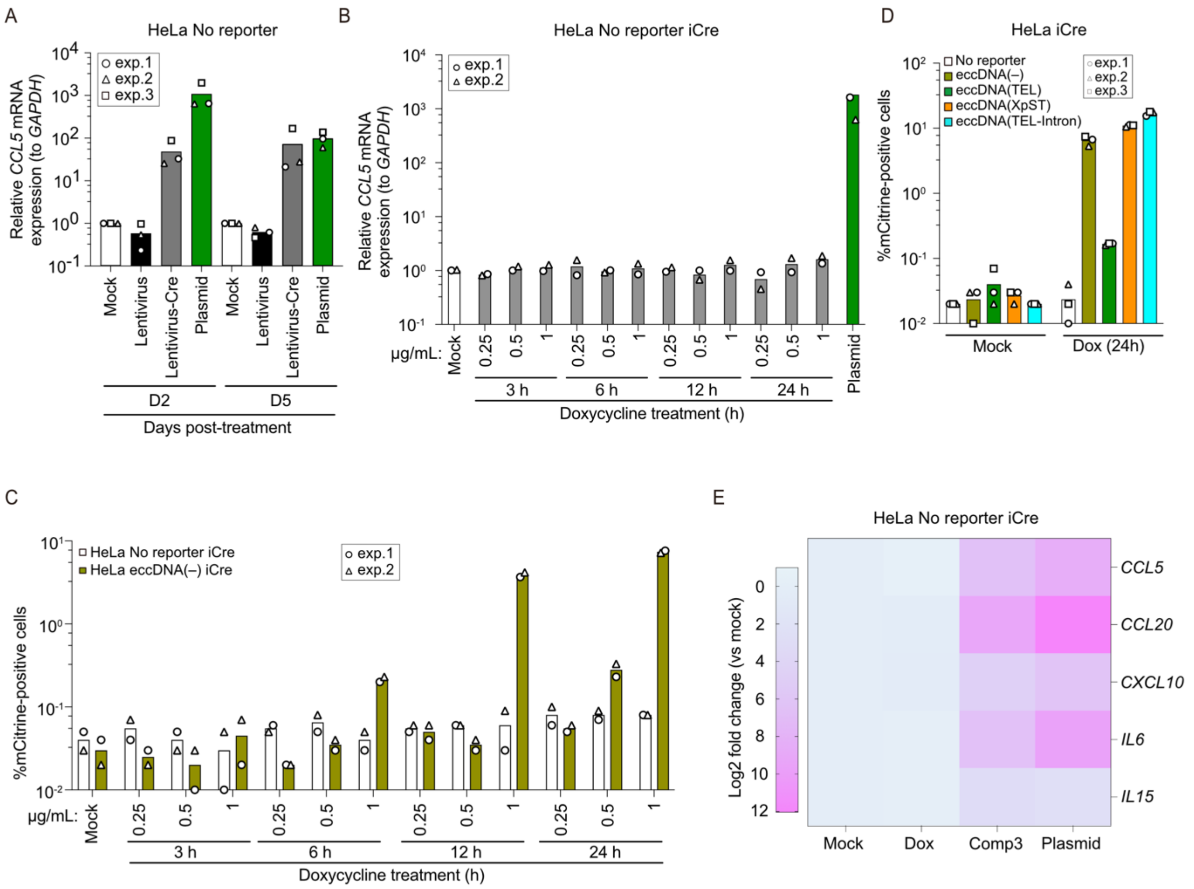


**Figure S4. Establishment and validation of dox-inducible Cre-dependent eccDNA reporter cells.** **(A)** Relative *CCL5* mRNA expression normalized to *GAPDH* in HeLa no reporter cells following mock treatment, lentiviral vector transduction, lentiviral Cre transduction, or plasmid transfection. RT-qPCR was performed on days 2 and 5 after the indicated treatment. **(B)** Relative *CCL5* mRNA expression normalized to *GAPDH* in HeLa no reporter iCre cells at the indicated dox concentrations and time points. Cells 24 hours after plasmid transfection served as a positive control. **(C)** Percentages of mCitrine-positive HeLa no reporter iCre and HeLa eccDNA(-) iCre cells at the indicated dox concentrations and time points. **(D)** Percentages of mCitrine-positive cells in the indicated HeLa eccDNA reporter iCre cells following mock or 24 h dox treatment. **(E)** Heatmap of relative cytokine and chemokine mRNA expression in HeLa no reporter iCre cells following mock treatment, 24 h dox treatment, Compound 3 (Comp 3) treatment, or plasmid transfection. Expression was normalized to *GAPDH*, and the color scale represents log2 fold changes relative to mock-treated cells. **Data information: (A-D)** Bars represent the mean of three independent (A, D) and two independent (B, C) biological replicates. **(E)** Heatmap values represent the mean of three independent biological replicates.

Figure S5: Validation of chromosome-derived eccDNA leakage into the cytosol

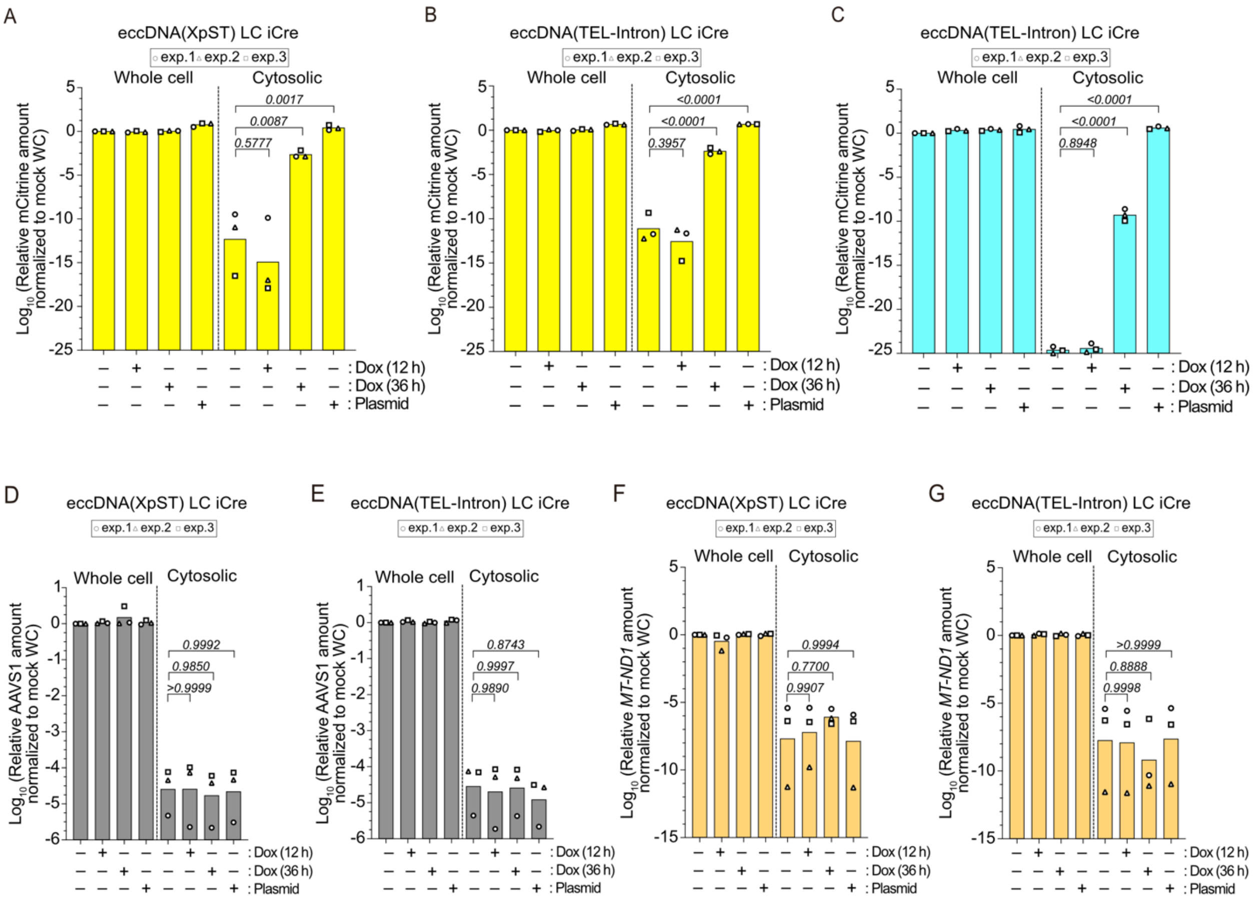


**Figure S5. Validation of chromosome-derived eccDNA leakage into the cytosol.**

**(A-G)** Relative mCitrine (A, B), telomere (C), AAVS1 (D, E), or *MT-ND1* (F, G) sequence amounts in whole-cell and cytosolic fractions of HeLa eccDNA(XpST) LC iCre (A, D, F) and eccDNA(TEL-Intron) LC iCre (B, C, E, G) mRuby3-STING cells, following dox treatment or plasmid transfection as shown in Fig. 4C. qPCR signals for each target were normalized to the corresponding mock whole-cell samples and shown as log10 fold change. **Data information:** Bars represent the mean of three independent biological replicates. Statistical significance was determined by ordinary one-way ANOVA followed by Dunnett's multiple-comparison test. P values are indicated for each comparison.

Figure S6: Chromosome-derived eccDNA induces modest STING accumulation

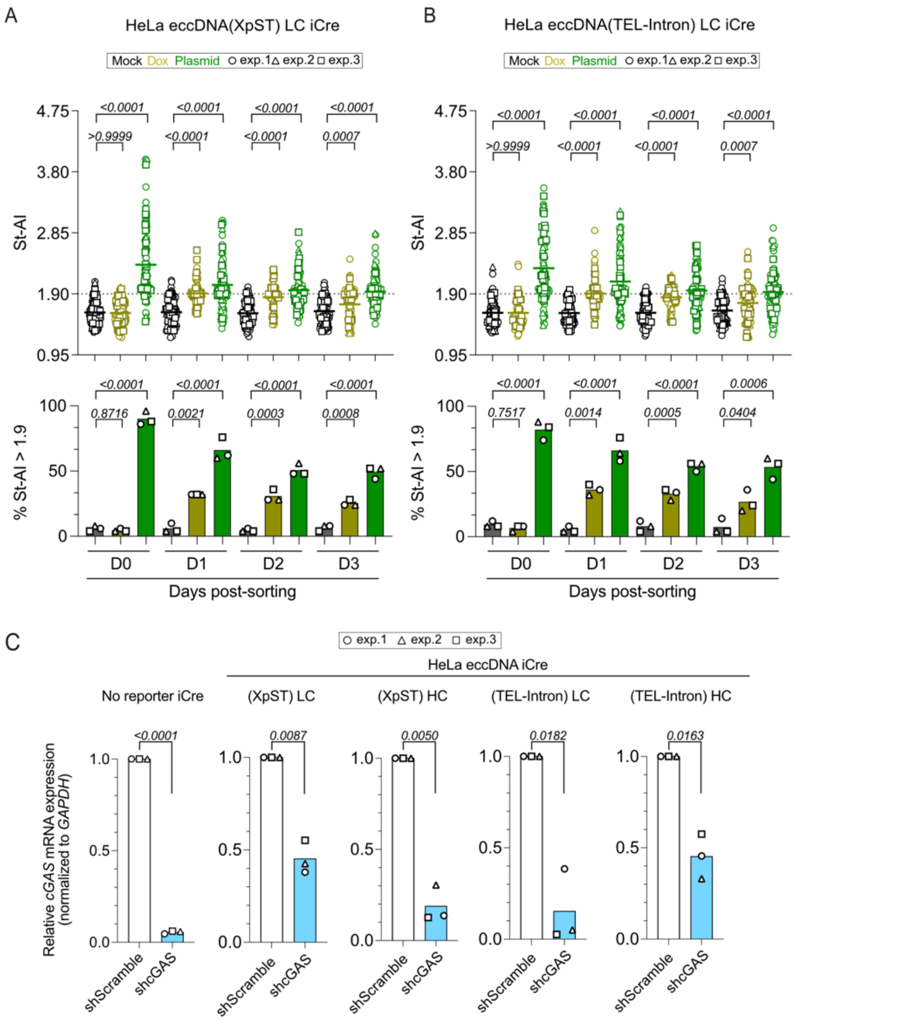


**Figure S6. Chromosome-derived eccDNA induces modest STING accumulation.**

**(A, B)** St-AI analysis of HeLa eccDNA(XpST) LC iCre (A) and eccDNA(TEL-Intron) LC iCre (B) mRuby3-STING cells following mock treatment, dox treatment, or plasmid transfection as shown in Fig. 5A. **(C)** RT-qPCR analysis of *cGAS* mRNA expression normalized to *GAPDH* in the indicated HeLa eccDNA reporter iCre mRuby3-STING cells following shScramble or shcGAS transduction as shown in Fig. 5F. **Data information: (A, B)** Bold lines represent the mean of three independent biological replicates (n = 100 cells in total, 50 cells in replicate 1 and 25 cells each in replicates 2 and 3). **(C)** Bars represent the mean of three independent biological replicates. Statistical significance in **(A, B)** was determined by the Kruskal-Wallis test followed by Dunn's multiple-comparison test (A, B: Top), by ordinary one-way ANOVA followed by Dunnett's multiple-comparison test (A, B: Bottom), and by Welch's t-test (C). P values are indicated for each comparison.

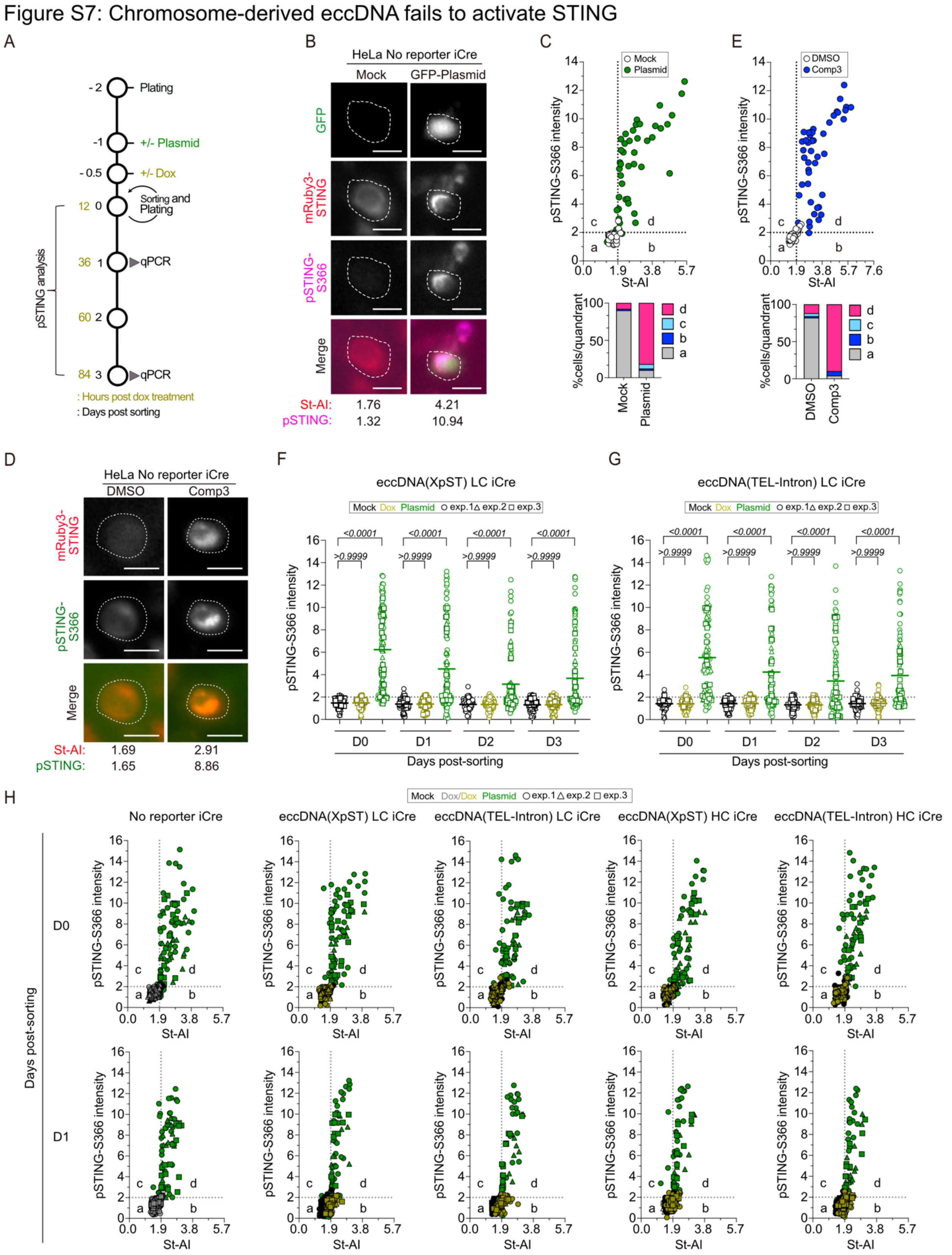


**Figure S7. Chromosome-derived eccDNA fails to activate STING.**

**(A)** Experimental workflow for pSTING analysis and RT-qPCR following sorting of mCitrine-positive eccDNA reporter cells or GFP-positive plasmid-transfected cells. **(B)** Representative images of GFP, mRuby3-STING, and pSTING-S366 IF in HeLa no reporter iCre mRuby3-STING cells following mock treatment or 24 h plasmid transfection. White dotted lines indicate cell boundaries.

Representative St-AI and pSTING-S366 values are shown below. Scale bars: 25 µm. **(C)** Top: single-cell pSTING-S366 intensity versus St-AI plots from cells in B. Bottom: percentages of HeLa no reporter iCre mRuby3-STING cells in each quadrant (a-d) defined by St-AI (threshold, 1.9) and pSTING-S366 (threshold, 2.0) on day 1 post-transfection. **(D)** Representative images of mRuby3-STING and pSTING-S366 IF in HeLa no reporter iCre mRuby3-STING cells following DMSO or 24 h Compound 3 (Comp 3) treatment, as shown in B. **(E)** Top: single-cell pSTING-S366 intensity versus St-AI plots from cells in D. Bottom: percentages of each quadrant are shown as in C. **(F, G)** pSTING-S366 intensity of HeLa eccDNA(XpST) LC iCre (F) and eccDNA(TEL-Intron) LC iCre (G) mRuby3-STING cells following the workflow shown in A. **(H)** Single-cell pSTING-S366 intensity versus St-AI plots of the indicated HeLa eccDNA reporter iCre mRuby3-STING cells on days 0 and 1 post-sorting. Percentages of each quadrant (a-d), defined as in C, are shown in Fig. 6E. **Data information: (F-H)** Data were obtained from three independent biological replicates (n = 100 cells in total, 50 cells in replicate 1 and 25 cells each in replicates 2 and 3). **(F, G)** Bold lines represent the mean. **(H)** Each dot represents a single cell. Statistical significance in (F, G) was determined by the Kruskal-Wallis test followed by Dunn's multiple-comparison test. P values are indicated for each comparison.

Figure S8: Validation of TREX1 knockdown

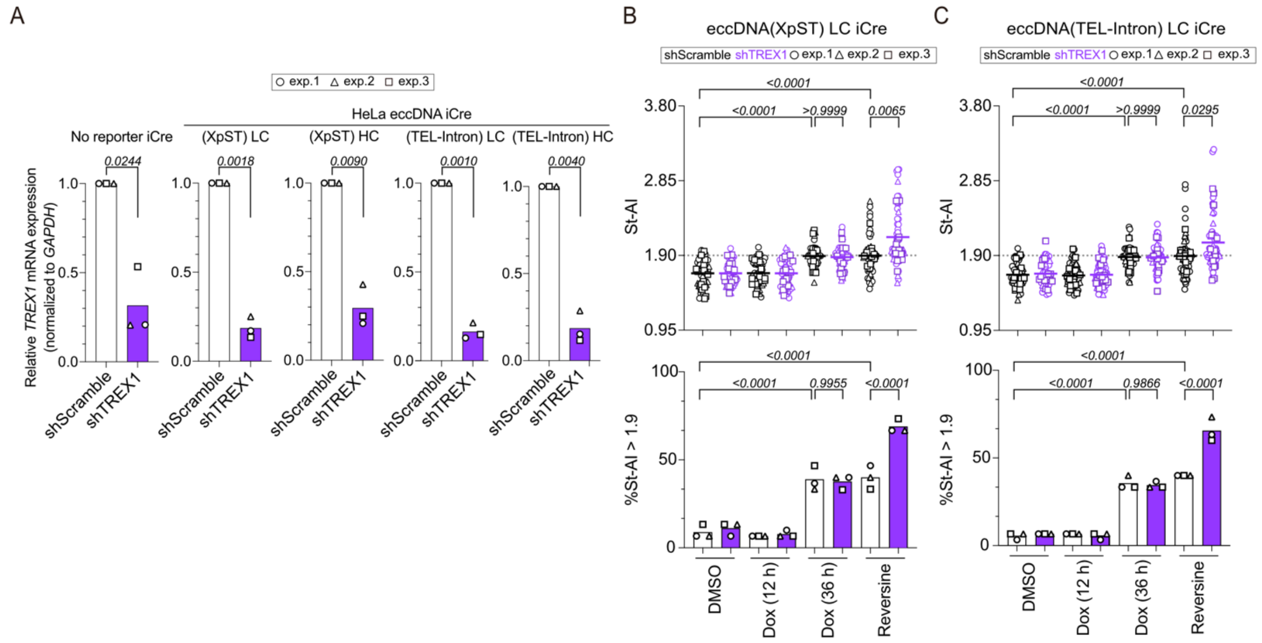


**Figure S8. Validation of TREX1 knockdown.**

**(A)** RT-qPCR analysis of *TREX1* mRNA expression normalized to *GAPDH* in the indicated HeLa iCre mRuby3-STING cell lines following shScramble or shTREX1 transduction, as shown in Fig. 7A. **(B, C)** Top: St-AI analysis of shScramble- or shTREX1-transduced HeLa eccDNA(XpST) LC iCre (B) and HeLa eccDNA(TEL-Intron) LC iCre (C) mRuby3-STING cells following DMSO treatment, 12 h and 36 h dox treatment, or 24 h reversine treatment. Bottom: the corresponding percentages of cells with St-AI > 1.9. For the 12 h dox condition, only mCitrine-positive cells were analyzed, whereas all cells were analyzed for the 36 h dox and reversine conditions. **Data information: (A)** Bars represent the mean. **(B, C)** Bold lines represent the mean. All data were obtained from three independent biological replicates (B, C, n = 60 cells in total, 30 cells in replicate 1 and 15 cells each in replicates 2 and 3). Statistical significance was determined by Welch's t-test (A), by the Kruskal-Wallis test followed by Dunn's multiple-comparison test (B, C: Top), and by ordinary one-way ANOVA followed by Šídák's multiple-comparison test (B, C: Bottom). P values are indicated for each comparison.
