## Supplementary material for "Uncoupling of STING Accumulation and Activation upon Chromosome-Derived Extrachromosomal Circular DNA Formation": Tables

**Table 1. List of Plasmids**

| Plasmid number (pMTH) | Name | Source |
|---|---|---|
| 127 | pLKO.1-shScramble | Addgene #1864 |
| 225 | pGFAP-Cre | Addgene #40591 |
| 295 | pBSII-NotI | This study |
| 380 | pMax-TurboGFP | (Sato & Hayashi, 2024) |
| 397 | pXp-franked sister | (Kagaya et al., 2020) |
| 688 | psPAX2 | Addgene #12260 |
| 689 | pCMV-VSV-G | Addgene #8454 |
| 753 | pCW-Cas9 | Addgene #50661 |
| 864 | pBS-sister(neo)-AAVS1 | (Kagaya et al., 2020) |
| 1050 | pLenti-bla-p2a-emiRFP703 | (Sato & Hayashi, 2024) |
| 1278 | pLentiCre-puro | This study |
| 1502 | pLKO.1-shcGAS-1 | (Sato & Hayashi, 2024) |
| 1633 | pCMV(CAT)T7-SB100 | Addgene #34879 |
| 1657 | pLKO.1-shTREX1-A | (Sato & Hayashi, 2024) |
| 1747 | pCm | This study |
| 1770 | pLenti-mRuby3-hSTING | This study |
| 1808 | pSV40_TelNN | (Huda et al., 2023) |
| 1824 | pBS-P-mCitC_N-P-neoR | This study |
| 1826 | pCWtetV16-Cre | This study |
| 1831 | pBS-P-mCitC_Esp3I-ClaI-mCitN-P-neoR | This study |
| 1853 | pBS-P-mCitC-TelNN691-N-P-neoR | This study |

| | | |
|---|---|---|
| 1864 | pBS-P-mCitC-XpSubTel691-N-P-neoR | This study |
| 1889 | pSBCm | This study |
| 1896 | pSBCm-P-mCitC_N-P-neoR | This study |
| 1898 | pSBCm-P-mCitC-telNN691_N-P-neoR | This study |
| 1899 | pSBCm-P-mCitC-Xpsubtel691_N-P-neoR | This study |
| 1930 | pBS-P-mCitC-N-SCBE-P-neoR | This study |
| 1942 | pBS-P-mCitC-TelNN691-2xNSE-mCitN-P-neoR | This study |
| 1987 | pBS-P-mCitC-N-TelNN691-P-neoR | This study |
| 1996 | pSBCm-P-mCitC-N-TelNN691-P-neoR | This study |

**Table 2. List of primers used for RCA, PCR and RT-qPCR**

| Primer number (MTH) | Name | 5'-3' Sequence | Purpose |
|---|---|---|---|
| 436 | mCitrine infusion-fwd | GTACCACAGGCAAACTTCCAGTTCCTTG | Genomic PCR |
| 477 | Neo seq rev | TCGACAAGACCGGCTTCCAT | Genomic PCR |
| 514 | NLS check R1 | GGCTGGACAGGCTAAGAAGA | RCA |
| 697 | TSK8C | CAGGGACCGGGACAAATAGAC | RCA |
| 1256 | mCitrine -fwd | TGGTAGTGATCCGCCAGTTG | Genomic qPCR |
| 1257 | mCitrine -rev | GCCGATAAGCAGAAGAACGG | Genomic qPCR |
| 1266 | AAVS1-arm L-fwd | ACAGCATGTTTGCTGCCTCCA | Genomic qPCR |
| 1267 | AAVS1-arm L-rev | AGTACCCAGAACCAGAGCCACA | Genomic qPCR |
| 1272 | KanR-seq-fwd | CGATGCCTGCTTGCCGAATATC | Genomic qPCR |
| 1273 | KanR-seq-rev | ATTCGCCGCCAAGCTCTTCA | Genomic qPCR |
| 1283 | No.114-IN-F | AGAAAGTATAGGAACTTCGCCTATCAA CAAGACATCAGTGGTTATGTTGAAAG | RCA |

| | | | |
|---|---|---|---|
| 1833 | oligo-dT | TTTTTTTTTTTTTTTT | cDNA (RT-PCR) |
| 1974 | GAPDH-fwd | CCTGCACCACCAACTGCTTA | RT-qPCR |
| 1975 | GAPDH-rev | GGCCATCCACAGTCTTCTGAG | RT-qPCR |
| 2264 | mCitrine-fwd | TTTGCCGCCAGAACACAGgatatcATGGTGTCTAAAGGTGAAGAG | Genomic qPCR |
| 2332 | cGAS-fwd | ATGCAAAGGAAGGAAATGGT | RT-qPCR |
| 2333 | cGAS-rev | TTTAAACAATCTTTCCTGCAACA | RT-qPCR |
| 2338 | cxcl10-fwd | GTGGCATTCAAGGAGTACCTC | RT-qPCR |
| 2339 | cxcl10-rev | GATGGCCTTCGATTCTGGATT | RT-qPCR |
| 2406 | il6-fwd | AGACAGCCACTCACCTCTTCAG | RT-qPCR |
| 2407 | il6-rev | TTCTGCCAGTGCCTCTTTGCTG | RT-qPCR |
| 2430 | MT-ND1-fwd | GGCTATATACAACTACGCAAAGGC | Genomic qPCR |
| 2431 | MT-ND1-rev | GGTAGATGTGGCGGGTTTTAGG | Genomic qPCR |
| 2461 | ccl20-fwd | AACCATGTGCTGTACCAAGAGT | RT-qPCR |

| | | | |
|---|---|---|---|
| 2462 | ccl20-rev | AAGTTGCTTGCTTCTGATTCGC | RT-qPCR |
| 2473 | ccl5-fwd | CCTGCTGCTTTGCCTACATTGC | RT-qPCR |
| 2474 | ccl5-rev | ACACACTTGGCGGTTCTTTCGG | RT-qPCR |
| 2477 | il15-fwd | AACAGAAGCCAACTGGGTGAATG | RT-qPCR |
| 2478 | il15-rev | CTCCAAGAGAAAGCACTTCATTGC | RT-qPCR |
| 2499 | TREX1-fwd | GCATCTGTCAGTGGAGACCACA | RT-qPCR |
| 2500 | TREX1-rev | CAGTGGTTGTGACAGCAGATGG | RT-qPCR |
| 3048 | 2S-XPFa infusion-rev | GCAGTTTCCCTGTGGTACAGATGAACTTCAGAG*T*G | RCA |
| 3066 | mCitrine-fwd | TGCTGTTGCCCGATAATCAC | Genomic qPCR |
| 3067 | mCitrine-rev | TTCCTGCTGCTGTGACAAAC | Genomic qPCR |
| 3279 | Telomere-fwd | CGGTTTGTTTGGGTTTGGGTTTGGGTTTGGGTTTGGGTT | Genomic qPCR |
| 3280 | Telomere-rev | GGCTTGCCTTACCCTTACCCTTACCCTTACCCTTACCCT | Genomic qPCR |

*Phosphorothioate modifications

**Table 3. List of antibodies**

| Antigen | Manufacturer | Catalog Number | Host Species | Dilution |
|---|---|---|---|---|
| Human pSTING-Ser366 | Cell Signaling Technology | 19781S | Rabbit | 1:200 |
| Alexa-488-conjugated goat anti-rabbit | Invitrogen | A11034 | Goat | 1:1000 |
| Cy5 labeled goat anti-rabbit IgG (H+L) | Cytiva | PA45004 | Goat | 1:500 |
